# Composition and stage dynamics of mitochondrial complexes in *Plasmodium falciparum*

**DOI:** 10.1101/2020.10.05.326496

**Authors:** Felix Evers, Alfredo Cabrera-Orefice, Dei M. Elurbe, Mariska Kea-te Lindert, Sylwia D. Boltryk, Till S. Voss, Martijn A. Huynen, Ulrich Brandt, Taco W.A. Kooij

**Affiliations:** Department of Medical Microbiology, Radboud Institute for Molecular Life Sciences, Radboud University Medical Center, PO Box 9101, 6500 HB Nijmegen, Netherlands; Radboud Institute for Molecular Life Sciences, Radboud University Medical Center, PO Box 9101, 6500 HB Nijmegen, The Netherlands; Centre for Molecular and Biomolecular Informatics, Radboud Institute for Molecular Life Sciences, Radboud University Medical Center, PO Box 9101, 6500 HB Nijmegen, the Netherlands; Electron Microscopy Center, RTC Microscopy, Radboud Institute of Molecular Life Sciences, Radboud University Medical Center, Geert Grooteplein 6525 GA Nijmegen, The Netherlands; Department of Cell Biology, Radboud Institute of Molecular Life Sciences, Radboud University Medical Center, Geert Grooteplein 6525 GA Nijmegen, The Netherlands; Department of Medical Parasitology and Infection Biology, Swiss Tropical and Public Health Institute, Basel, Switzerland; University of Basel, Basel, Switzerland; Cologne Excellence Cluster on Cellular Stress Responses in Aging-Associated Diseases (CECAD), University of Cologne, Cologne, Germany

## Abstract

Our current understanding of mitochondrial functioning is largely restricted to traditional model organisms, which only represent a fraction of eukaryotic diversity. The unusual mitochondrion of malaria parasites is a validated drug target but remains poorly understood. Here, we apply complexome profiling to map the inventory of protein complexes across the pathogenic asexual blood stages and the transmissible gametocyte stages of *Plasmodium falciparum*. We identify remarkably divergent composition and clade-specific additions of all respiratory chain complexes. Furthermore, we show that respiratory chain complex components and linked metabolic pathways are up to 40-fold more prevalent in gametocytes, while glycolytic enzymes are substantially reduced. Underlining this functional switch, we find that cristae are exclusively present in gametocytes. Leveraging these divergent properties and stage dynamics for drug development presents an attractive opportunity to discover novel classes of antimalarials and increase our repertoire of gametocytocidal drugs.

## Introduction

Malaria parasites harbour only a single, indispensable mitochondrion with a minimalistic mitochondrial DNA (mtDNA) encoding just three proteins: COX1, COX3, and CYTB(1, 2). The latter is the target of the potent antimalarial atovaquone (3). Its activity on asexual blood-stage (ABS) parasites is not directly mediated by inhibition of the oxidative phosphorylation (OXPHOS) pathway but by blocking ubiquinone regeneration required to sustain *de novo* pyrimidine biosynthesis (4). *Plasmodium* gametocyte development and mosquito colonization on the other hand are critically dependent on multiple mitochondrial functions including an active respiration (5-8). Another remarkable feature observed in the murine malaria model parasite *Plasmodium berghei*, is the apparent absence of cristae in ABS (7). *P. berghei* gametocytes, however, do possess these inner mitochondrial membrane folds where OXPHOS complexes typically accumulate. Due to high sequence diversity and the poor mitochondrial targeting predictions, much of the *Plasmodium* mitochondrial proteome remains undisclosed. This is painfully illustrated by our limited understanding of a pathway as central to mitochondrial functioning as the OXPHOS pathway. The marked absence of complex I is compensated by a single subunit type II NADH:ubiquinone oxidoreductase. For CII-V, only 23 likely orthologues of 48 canonical components have been identified (9) and no comprehensive biochemical analysis of any of the individual complexes has been done so far. Recent studies in the related apicomplexan parasite *Toxoplasma gondii* have suggested a divergent and unusual composition of cytochrome *c* oxidase (CIV; (10) and F_1_F_O_-ATP synthase (CV)(11, 12). To this date, only limited data about multiprotein complexes in *Plasmodium* species are available. Function annotations, co-expression patterns, and homology data have been integrated *in silico* to predict possible protein interactions (13, 14). However, this approach is hampered by the lack of annotated orthologues for many proteins, limited temporal resolution of expression data, and imperfect correlation between transcription and translation timing in *Plasmodium* species (15). A systematic yeast two-hybrid screen generated protein interaction data for 25% of the proteome (16) but pairwise expression of protein fragments outside of their native context is not necessarily representative or suitable to uncover all relevant protein-protein interactions, especially in the case of multiprotein complexes.

Recent progress in label-free quantitative mass spectrometry (MS) combined with very high sensitivity and speed microscale native fractionation techniques offers the prospect to uncover protein associations by comigration or co-elution (17). Blue native polyacrylamide gel electrophoresis (BN-PAGE;(18) offers high resolution separation of intact complexes over a wide mass range without requiring genetic interventions or prior modifications of the sample (17). This approach, termed complexome profiling, provides the inventory of protein complexes in a single experiment. It has allowed for major advances by finding novel components of OXPHOS complexes and uncovering assembly intermediates and interactions in human (19), plant (20) and yeast (21) mitochondria. A prior study by Hillier *et al*. (22) has demonstrated that this approach is feasible and effective in *Plasmodium* spp., but due to different scope, sample complexity and comparatively harsh detergent conditions has failed to identify assembled OXPHOS complexes.

Here, we apply complexome profiling to preparations of *Plasmodium falciparum* ABS parasites and gametocytes enriched for mitochondria. The extensive datasets provide a wealth of information on *Plasmodium* protein complexes allowing the identification of numerous previously suggested and novel components of all OXPHOS complexes. Critically, we uncover stark OXPHOS complex abundance differences between asexual and sexual blood-stage parasites, consistent with the metabolic switch hypothesis and coinciding with the appearance of cristae as supported by our ultrastructural observations. Further analysis of these parasite-specific OXPHOS components could pave the way towards novel drug targets and enables a better understanding of this divergent and fascinating mitochondrial biology.

## Results

### Stage-specific mitochondrial ultrastructure in *P. falciparum*

To provide ultrastructural support for the increasing evidence for stage-specific mitochondrial metabolism and function, we performed transmission electron microscopy (TEM) of NF54 wildtype *P. falciparum* ABS parasites and stage V gametocytes (Fig. 1). Prior TEM-based investigations have demonstrated the presence of cristae in *P. berghei* gametocytes while ABS parasites were acristate (7). Cristae in *P. falciparum* stage IV gametocytes have also been suggested, but low image quality and absence of ABS micrographs, do not allow definitive conclusions (23). Our data confirm the stage-specific presence of cristae inside the *P. falciparum* mitochondrion (Fig. 1). In ring-stage parasites, the mitochondrion is elongated, acristate and not very electron dense. While significantly larger, the overall appearance is unchanged in trophozoites. In mature schizonts, each daughter merozoite harbours one small, acristate, electron-lucent mitochondrion in close proximity to one four-membrane-bound apicoplast. We also observed that these organelle pairs are distributed to merozoite compartments prior to inclusion of the nuclei (Supplementary Fig. 1). In gametocytes, clear and abundant internal membranous structures are observed within the mitochondrion, which we assume to be tubular cristae due to their resemblance to tubular cristae observed in steroid-producing (24) cells and *T. gondii* (25). Additionally, the organelle appears more electron-dense than in ABS and covers larger distinct areas suggesting an increase in size and level of branching (Supplementary Fig. 2). The multiple mitochondrial sections without apparent connection are assumed to be part of one heavily branched mitochondrion (26) but appear to be distinct as the 3D conformation cannot be appreciated in the 2D micrographs. This obvious discrepancy between all ABS parasites on one side and mature gametocytes on the other side prompted us to investigate how these changes are reflected at the protein level.

**Figure 1.**
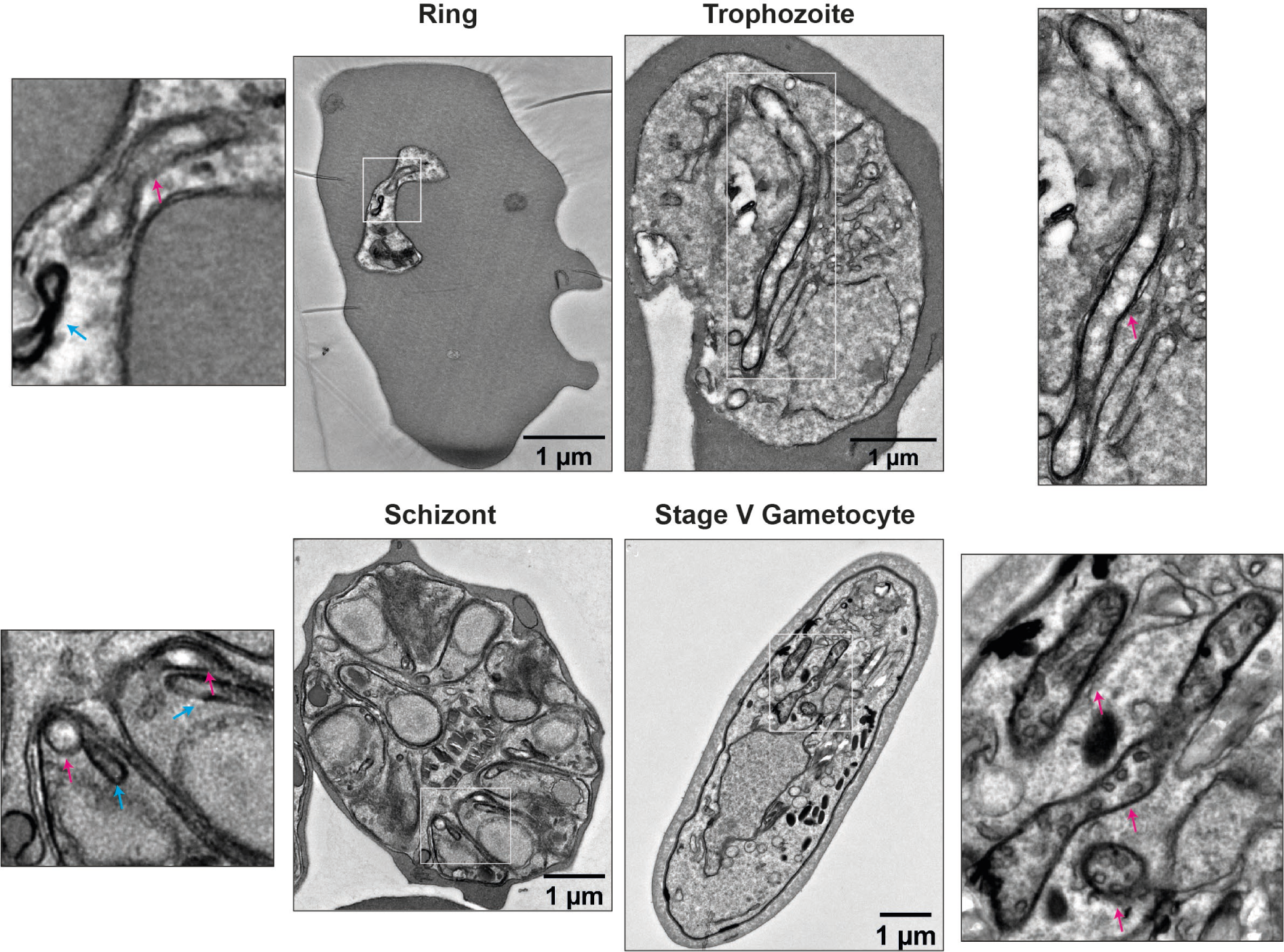
Representative electron micrographs of *Plasmodium falciparum* blood stages. Enlarged sections show ultrastructural differences between the mitochondrion (red arrow) in ABS parasites and gametocytes. The mitochondrion presents as an electron-lucent, acristate structure during ABS development, while the gametocyte mitochondrion appears electron-dense and packed with tubular cristae. Also note the close proximity of the four-membrane-bound apicoplast (green arrow) in ring- and schizont-stage parasites.

### Complexome profiling of *Plasmodium falciparum*

Migration patterns of individual proteins were obtained by complexome profiling of mixed ABS parasites and stage V gametocytes (GCT). We employed three different enrichment methods (1: syringe lysis with saponin; 2: syringe lysis without saponin; 3: nitrogen cavitation), two different detergents (D: digitonin; M: n-Dodecyl β-D-maltoside, DDM), as well as two genetic backgrounds were analysed (see Methods and Supplementary Table 1 for further details). Samples were named according to the combination of these three parameters and lettering indicating different replicates (Supplementary Table 1). Saponin treatment during shearing was used to test whether specific depletion of relatively cholesterol-rich non-mitochondrial membranes by saponin (27) could improve mitochondrial enrichment. With some notable exceptions, which will be described later, the results were consistent across all enrichment methods. Using the stronger detergent DDM did not lead to detection of more proteins or assembled complexes (Supplementary Table 1) and was consequently omitted in favour of digitonin solubilization for gametocyte samples. To overcome the challenge of obtaining sufficient gametocyte material, we used a recently established inducible gametocyte producer line (NF54/iGP2) (Boltryk et al., unpublished) that allows for the synchronous mass production of gametocytes through conditional overexpression of gametocyte development 1 (GDV1), an activator of sexual commitment (28). We observed no marked differences in proteome or morphology for ABS and GCT stages between the initial samples prepared with wild-type NF54 (GCT1Da, ABS1Ma, ABS1Da) and all remaining samples prepared with NF54/iGP2 parasites (Supplementary Table 1). Across all samples (662 fractions) a total of 1,759 unique proteins were identified (Supplementary Information S1). All raw and processed data generated in this study was deposited at the ComplexomE profiling DAta Resource (CEDAR; www3.cmbi.umcn.nl/cedar/browse/experiments/CRX23). It should be noted that abundant proteins from other *P. falciparum* cell compartments were also readily identified. Taking advantage of the latter and to validate the approach, we first investigated whether well-known and previously described complexes could be identified correctly and what the impact was of different isolation methods (Fig. 2).

**Figure 2.**
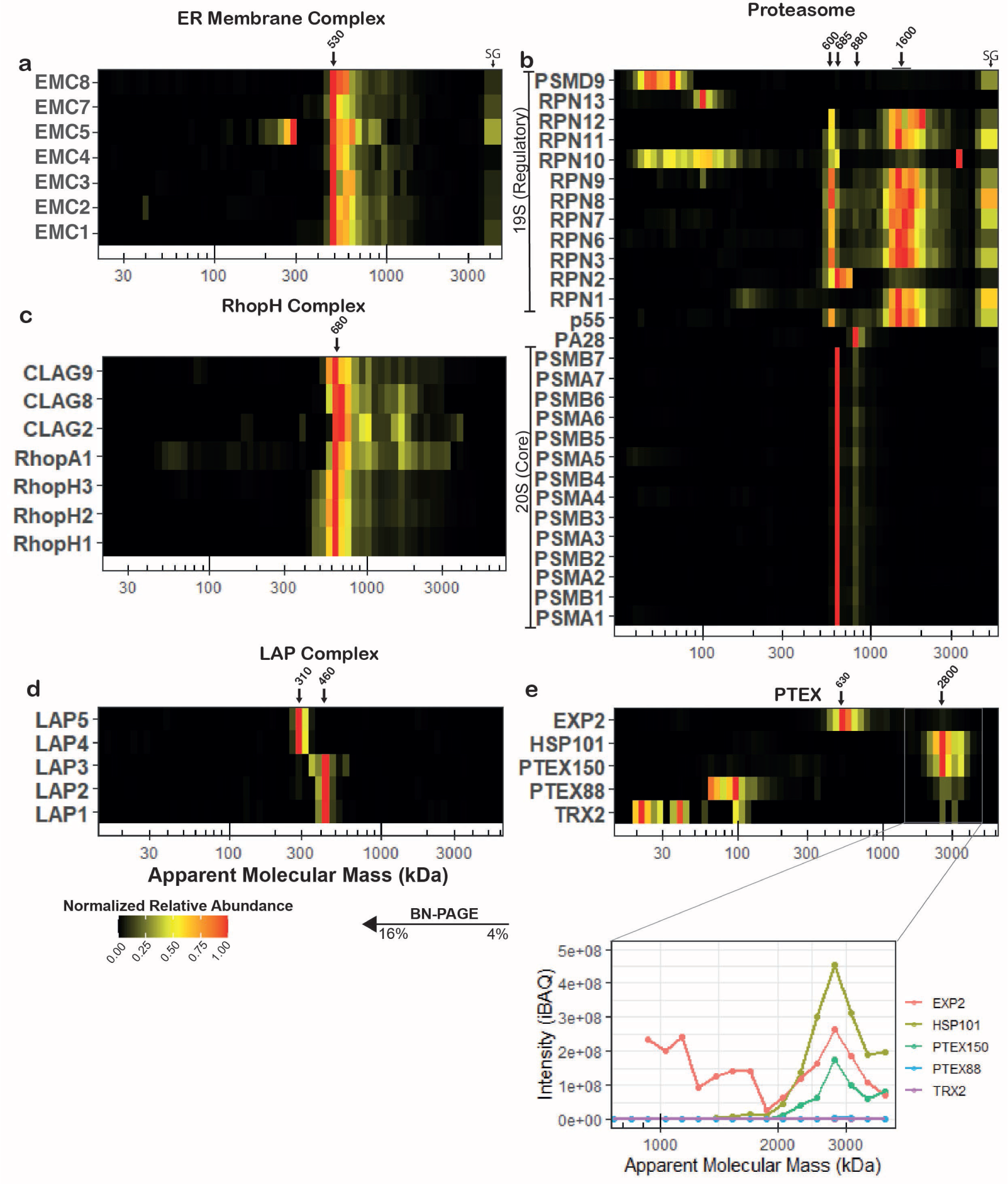
Migration profiles of proteins associated with previously described complexes. (**a**) Putative EMC components, representative heat map from sample ABS3D. All detected EMC components comigrate at an M_r_^app.^ of ∼530 kDa. (**b**) Proteasome components, representative heat map from sample ABS3D. 20S components comigrate at an M_r_^app.^ of ∼690 kDa and to a lesser degree at ∼880 kDa. 19S regulatory components comigrate at two distinct sizes, a major proportion at ∼1600 kDa and a smaller fraction at ∼600 kDa. (**c**) RhopH complex, representative heat map from sample ABS1Da. PF3D7_0220200 (RhopHA1) shared RhopH complex pattern consistently and thus was putatively assigned to the RhopH complex. (**d**) LAP complex, representative heat map from sample GCT1Da. LAP complex components in gametocytes migrated as two distinct subcomplex consisting of LAP1-3 (∼460 kDa) and LAP4-5 (∼310 kDa) respectively. Additionally, a faint putative assembly intermediate consisting of LAP2 and LAP3 was observed at ∼310 kDa. (**e**) PTEX, representative heat map (upper panel) and line chart of iBAQ values in the 750-4000 kDa mass range from sample ABS1Da. The PTEX complex including auxiliary subunits can be observed to comigrate at an M_r_^app.^ of ∼2.8 MDa. Due to high proportion of EXP2 present as the homooligomeric EXP2 complex at ∼800 kDa, membership is only evident when comparing absolute intensity values (lower panel) instead of normalized abundances (heat map). PTEX88 and TRX2 have much lower intensities in this mass range than core components. SG, stacking gel. Corresponding Gene IDs can be found in Supplementary Table S1

### Validation with common eukaryotic and *Plasmodium*-specific complexes

The protein folding/degradation-involved endoplasmic reticulum membrane complex (EMC) genes are conserved across most eukaryotes (29). We detected comigration of all putative EMC components with the exception of EMC6 at an apparent molecular mass (M_r_^app.^) of ∼550 kDa (Fig. 2a). Previous proteomics experiments also failed to detect EMC6 in ABS parasites and gametocytes (30) despite having similar transcription profiles as other EMC components (https://plasmodb.org), indicating challenging detection by MS or absence. The M_r_^app.^ differs from the predicted mass of 294 kDa, possibly due to presence of N-glycans on multiple copies of some components (31, 32), interaction with other unidentified proteins (33), or a larger oligomeric state. Nevertheless, consistent comigration of the identified subunits provides strong evidence for a canonical EMC assembly in *P. falciparum*.

The presence of a proteasome complex represents another universal eukaryotic feature. A prior study has elucidated the composition and M_r_^app.^ of the *P. falciparum* 20S proteasome (34). We confirmed these results, identifying all 14 subunits of the 20S proteasome comigrating as a clearly defined complex at ∼690 kDa and a less abundant, slightly larger assembly (Fig. 2b). Most components of the regulatory 19S particle comigrated in a dominant large and secondary small assembly. The lack of comigration between regulatory components and the core 20S proteasome suggests limited stability of the 26S or 30S assemblies under these conditions. Interestingly, the regulatory subunit 2 (RPN2) seemed to associate with both the dominant 20S and 19S assemblies, while the proteasome activator subunit 28 (PA28) was found exclusively associated with the larger 20S complex. The putative 26S proteasome non-ATPase regulatory subunit 9 (PSMD9) and regulatory subunit 13 (RPN13) were not found comigrating with any of the observed assemblies. We observed that saponin treatment depletes all proteasome-associated assemblies from the respective profiles, suggesting either reduced cytosolic contaminants or a specific detergent-complex interaction upon saponin treatment (Supplementary Fig. 3). Conversely, the studied membrane proteins were not significantly affected under these conditions.

To facilitate waste removal and nutrient uptake through their host cell membrane, malaria parasites have evolved a unique complex composed of the high molecular mass rhoptry proteins (RhopH;(35). Initially thought to be composed of three subunits, recent studies have implicated additional proteins and estimated its mass as ∼670 kDa (36). Although CLAG3.2 was not detected in any of the samples, presumably due to a high overlap in shared and consequently non-unique peptides with RhopH1, our observations otherwise confirmed the composition and size of this extended RhopH complex and suggested a new component, which we termed RhopH associated protein 1 (RhopA1; PF3D7_0220200; Fig. 2c). RhopHA1 comigrated consistently with the previously established components and at the predicted mass under all conditions. It has two predicted transmembrane helices, a PEXEL motif, and no detectable homologues outside the *Plasmodium* genus.

Successful development in the mosquito vector requires expression of six LCCL domain-containing proteins that form a complex in the crystalloid, an organelle unique to *Plasmodium* insect stages (37, 38). In *P. berghei*, early assembly of this complex is prevented through translational repression of LAP4-6 in gametocytes (39). In *P. falciparum*, LAP1-5 are transcribed from stage II gametocytes onwards and are readily detected by MS, while there is no MS evidence of LAP6 in gametocytes and transcription only commences in stage V gametocytes (40, 41). Immunofluorescence microscopy with one antiserum suggested the presence of LAP6 but at an entirely different localization than the other LAP proteins (38, 42). We did not detect LAP6 in any of the samples but found that in stage V gametocytes LAP1-3 and LAP4-5 formed two distinct subcomplexes (Fig. 2d). We also identified a likely LAP2-3 assembly intermediate. The apparent masses of the complexes resemble the sum of their constituents, suggesting that LAP6 was indeed not present. An interesting explanation could be that instead of repressing translation of LAP4-6 as in *P. berghei*, only LAP6 is repressed in *P. falciparum*, which then functions as the assembly factor bringing the two subcomplexes together after fertilization.

The *Plasmodium* translocon of exported proteins (PTEX) is a malaria parasite-specific protein complex essential for the export of parasite proteins into the host erythrocyte(43, 44). The structure of the core complex consisting of three proteins, EXP2, PTEX150, and HSP101, is now well established(45), while function and even association of the non-essential auxiliary subunits PTEX88 and TRX2 remains less clear(46). Our data indicated that the majority of EXP2 is present independently from the PTEX (Fig. 2e), consistent with its second proposed function as a homooligomeric nutrient channel (47). Reassuringly, when comparing intensity based absolute quantification (iBAQ) values, the three core components showed comparable intensities (lower panel Fig. 2e). Conversely, the majority of PTEX88 and TRX2 migrated at the predicted size of the monomer, while low intensity comigration with the core complex was inconsistent across the samples and even replicates, suggesting their presence in only a subset of complexes or a limited association under the experimental conditions used (Supplementary Fig. 4). The fact that complexome profiling helps to distinguish the presence of proteins in different (sub)assemblies, highlights a key advantage over more conventional approaches to investigate interactions of promiscuous components or assess assembly pathways.

### Divergent composition and abundance dynamics of CIII and CIV

Having validated our complexome profiling approach, we next focussed on the OXPHOS complexes. All canonical components of cytochrome *bc*_*1*_ (CIII) with obvious *Plasmodium* orthologues, *i*.*e*. CYTB, CYTC1, the Rieske subunit, QCR7, and QCR9, comigrated (Fig. 3a), with the notable exception of *Pf*QCR6 (PF3D7_1426900), that was not detected potentially due to its small size, hydrophobicity, and limited generation of identifiable unique peptides. As observed in plants (48), MPPα and MPPβ also associated with CIII coupling processing peptidase activity to a structural role in replacing the so called core proteins. Four additional proteins comigrated consistently with CIII subunits. We identified PF3D7_0306000 as a likely orthologue of QCR8 (E=6×10^−6^), while the other three proteins, which we termed respiratory chain complex 3 associated proteins 1-3 (C3AP1-3; Table 1), were found almost exclusively in Apicomplexa and lack any detectable sequence homology with characterized proteins (Fig. 4a). In other species, CIII forms a dimer of 470-500 kDa (20, 49). At ∼730 kDa, the M_r_^app.^ of *P. falciparum* CIII was considerably larger but similar to the 690 kDa expected molecular mass for an obligatory dimer including the newly identified subunits.

**Table 1.**
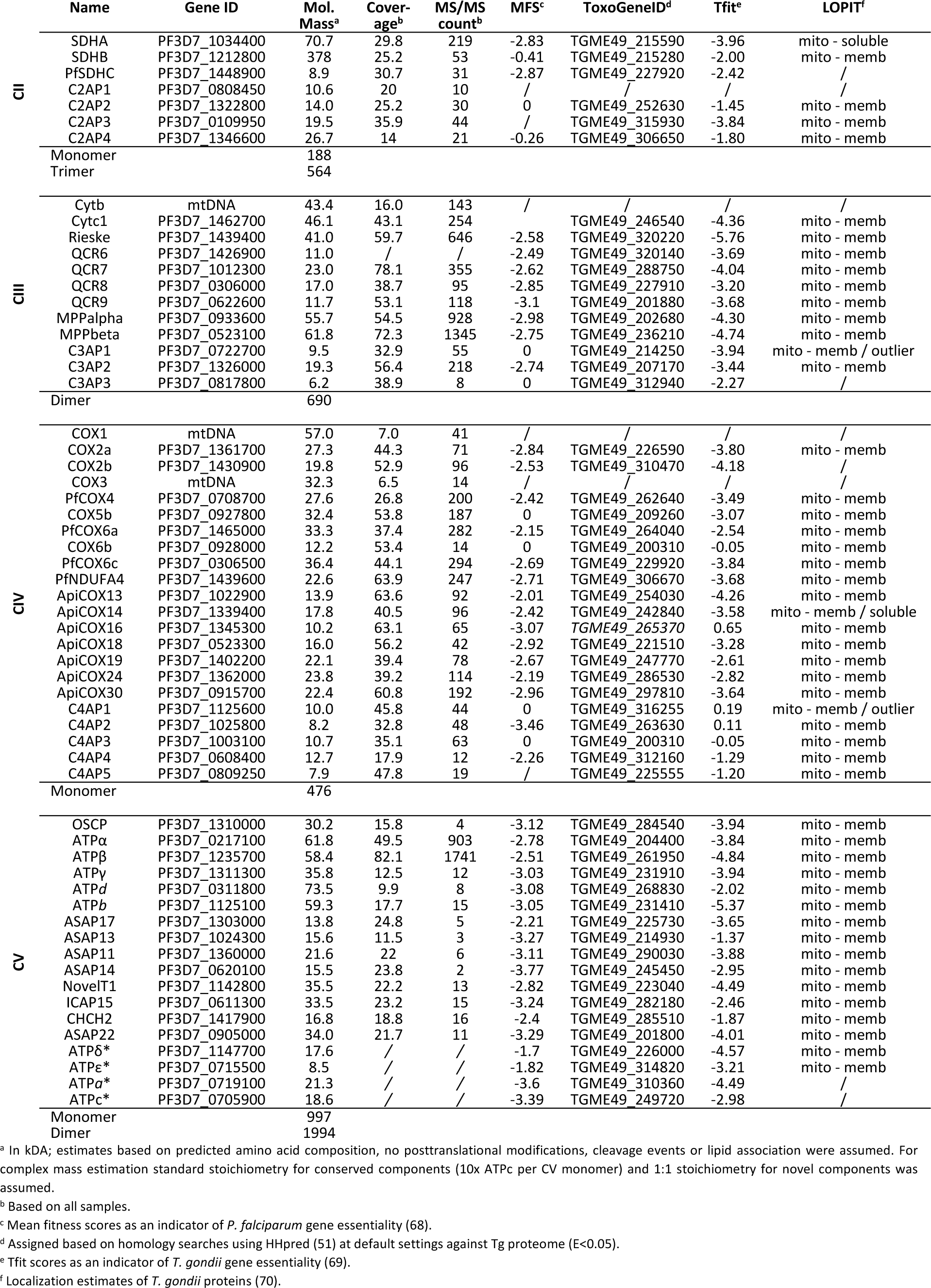
Divergent composition of the respiratory chain complexes in *Plasmodium falciparum*.

**Figure 3.**
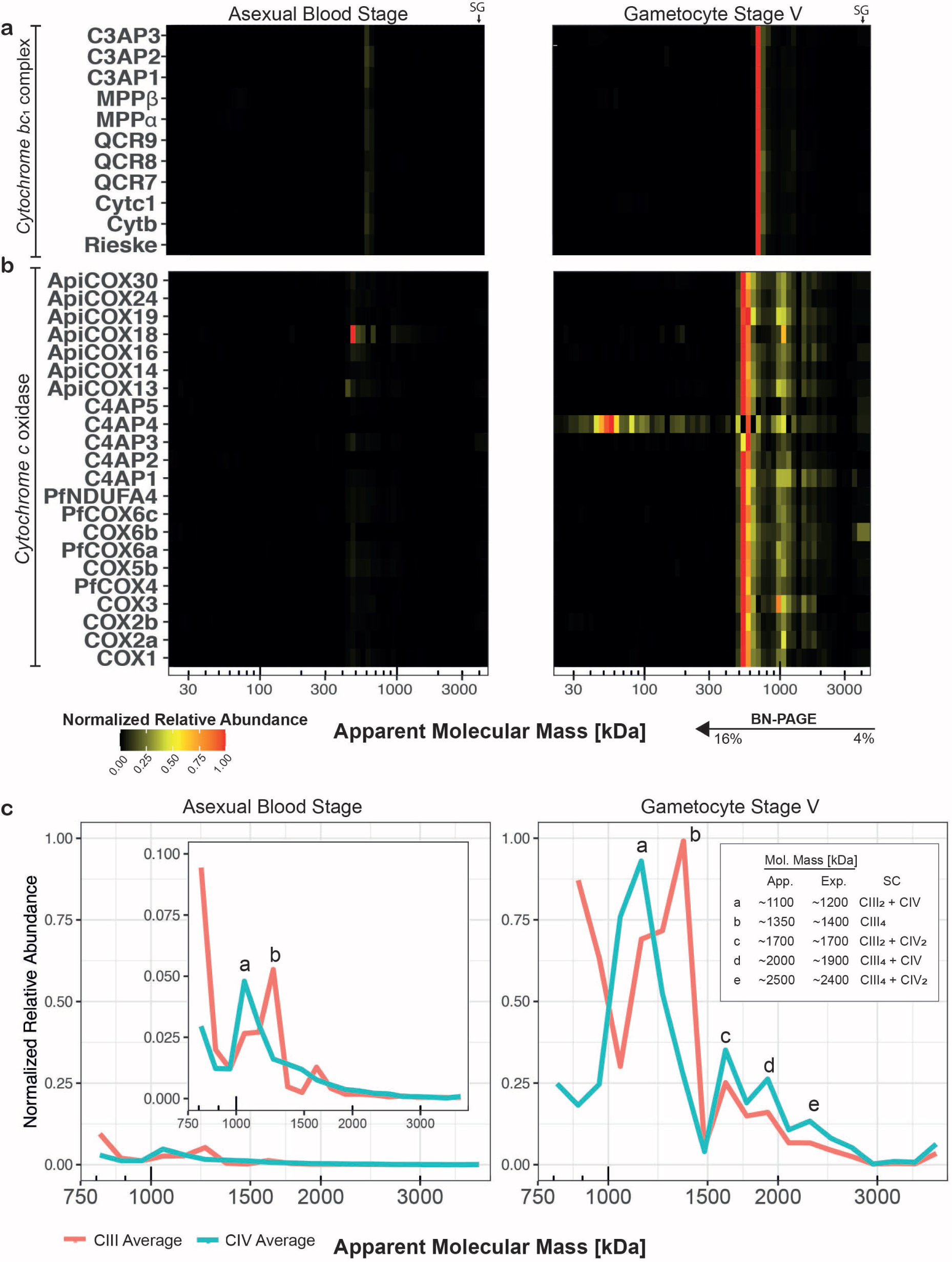
Migration and relative abundance of canonical and putatively associated components of respiratory chain complexes III and IV. An abundance of 1 (red) represents the highest iBAQ value for a given protein between both samples. (**a**) Heatmap showing comigration of canonical CIII components as well as putative novel components migrating at an M_r_^app.^ of ∼730 kDa respectively in ABS parasites (left) and gametocytes (right). (**b**) Heatmap showing comigration of canonical CIV components as well as putative novel components migrating at an M_r_^app.^ of ∼570 kDa as well as relative abundance in ABS parasites (left) and gametocytes (right). (c) For detailed analysis of higher-order assemblies, intensity values at M_r_^app.^. >700 kDa were renormalized and visualized in a line plot. Different putative supercomplexes were observed in ABS and gametocytes, denoted with lettering and described in graph inlet. apparent, approximate molecular mass based on migration profile; expected, expected molecular mass based on composition observed in this study; SC, supercomplex; CIII_2_, obligatory CIII dimer; CIII_4_, association of two CIII dimers; CIV, CIV monomer; CIV_2_, CIV dimer.

**Figure 4.**
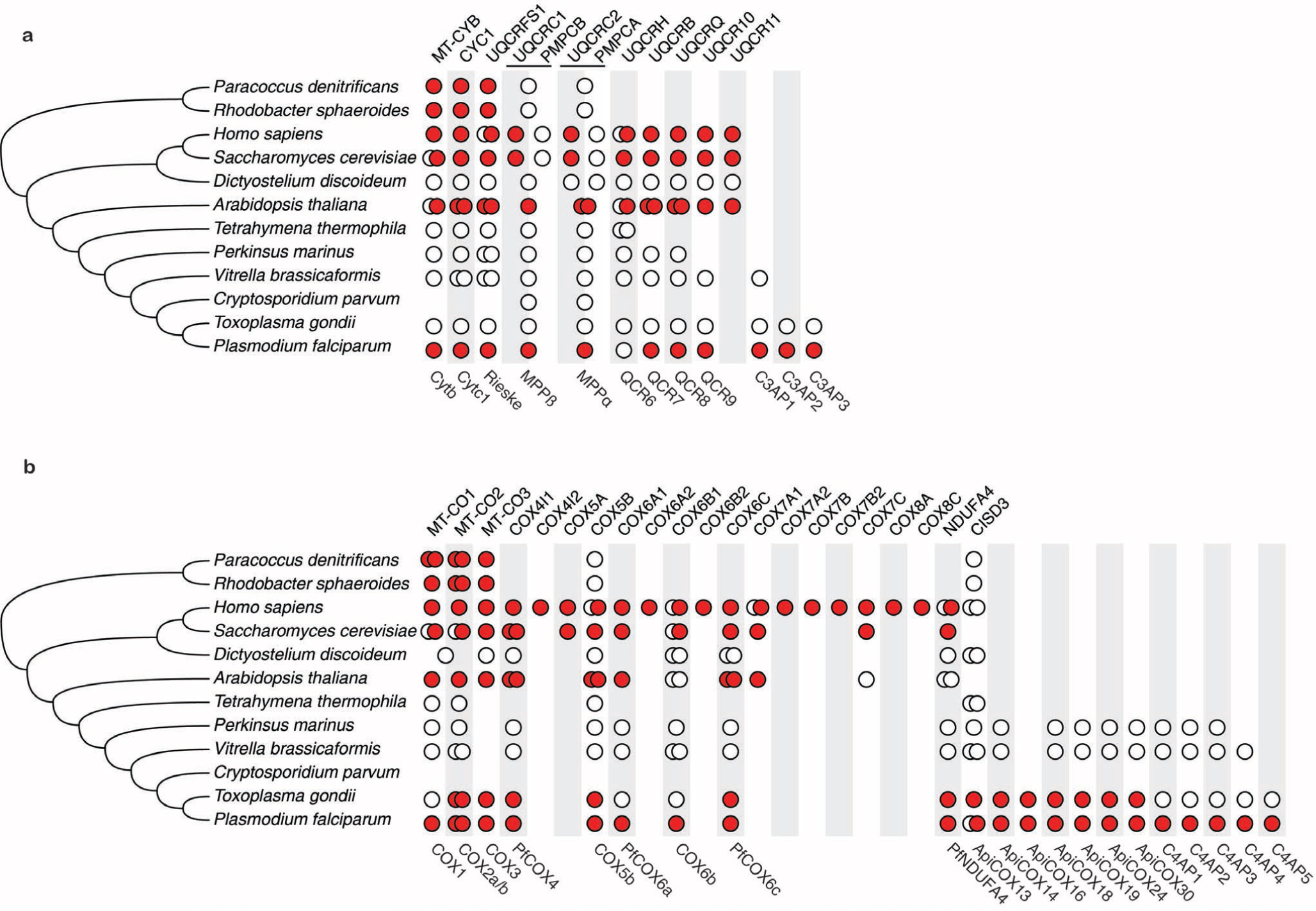
Evolution of complex III (A) and complex IV (B) subunit composition in model species and in the lineage leading to *P. falciparum*. The composition of each complex (rows) is based on data from model species and proteins from *P. falciparum* found to comigrate with that complex in this study. Colours depict levels of evidence (red, experimental evidence; white, genomic evidence) linking the subunit to the enzyme. Double circles represent presence of paralogs and their colour indicates experimental evidence linking them to the complex. For PMPCA/UQCRC2 and PMPCB/UQCRC1, in cases where there has not been a gene duplication, the protein is indicated in the middle between the two columns Human gene symbols are shown on top; *P. falciparum* gene names are shown below the conservation matrix.

So far, only five canonical subunits of *Plasmodium* cytochrome *c* oxidase (CIV) have been identified, *i*.*e*. COX1, COX2, COX3, COX5b, and COX6b. COX2 is generally encoded in the mtDNA but in Apicomplexa and Chlorophyceae the gene has been split in two and relocalized to the nucleus (50). The resulting protein fragments, COX2a and COX2b, were both retrieved in the complexome profiles. Recent research has shown a highly divergent composition of CIV in *T. gondii*, containing 11 subunits specific to Apicomplexa (10). Comigration of orthologues of all of these subunits with canonical CIV components confirmed this atypical CIV composition for *P. falciparum* (Fig. 3a). Three proteins that were deemed apicomplexan-specific by Seidi *et al*. (10) have significant sequence similarity to canonical CIV subunits (E<0.01; Fig. 4b), *i*.*e. Pf*COX6A (PF3D7_1465000), *Pf*NDUFA4 (PF3D7_1439600), and *Pf*COX4 (PF3D7_0708700). Furthermore, we identified a minimal level (E= 0.74) of sequence conservation at the C-terminus of PF3D7_0306500 with COX5C in *Arabidopsis thaliana*, which is orthologous to COX6C from Metazoa and COX9 in fungi. In support of its potential orthology, the conserved residues are all located at the interface of the transmembrane region of COX9 and the other CIV subunits (Supplementary Fig. 5). Through sequence profile-based searches the comigrating protein PF3D7_1345300, was identified as orthologous to ApiCOX16 (E=4.5×10^−41^), which was previously assumed to be *T. gondii* specific. In addition, we identified five uncharacterized, largely myzozoan-specific proteins that consistently comigrated with the complex and which we termed respiratory chain complex 4 associated proteins 1-5 (C4AP1-5; Table 1).

The complexome profiles suggested staggering abundance differences between ABS parasites and stage V gametocytes (Fig. 3a). Following enrichment by nitrogen cavitation, intensity values for suggested CIII and CIV components are on average 9-fold and 20-fold higher in gametocytes than in ABS parasites. This is not contradicted by the seemingly high relative abundance of ApiCOX18 in the ABS heatmap, since in this case the much lower abundance approached the detection level causing a normalization artifact. When averaging all digitonin-solubilized samples, stage-differences were 6-fold and 23-fold for CIII and CIV components, respectively (Supplementary Table 2).

Finally, to better visualize the presence of higher-order assemblies, we renormalized relative abundances based on intensity values detected at an M_r_^app.^ >750 kDa. Thus, we identified complex-specific higher-order assemblies possibly corresponding to the CIII dimer associating with an CIV monomer (Fig. 3b, peak a) and dimer (peak c). We also observed putative association of two CIII dimers (peak b) and of two CIII dimers with an CIV monomer (peak d) and dimer (peak e). It is noteworthy, that the latter larger putative respiratory supercomplexes peaks are exclusively observed in the gametocyte samples. Even when disregarding absence of distinct peaks, relative intensity at the supercomplex sizes compared to the dominant CIII/CIV bands in ABS parasites, is very low compared to gametocyes (Supplementary Table 2). This could either be caused by their absence from ABS parasites or falling below the detection threshold due to overall lower abundance of OXPHOS complex.

### Composition of respiratory chain complexes III and IV in an evolutionary context

To examine the evolution of *P. falciparum* CIII and CIV in detail, we mapped both the gains and the losses of their respective subunits along an evolutionary tree (Fig. 4). To ensure maximum sensitivity, homology detection was done using HHpred (51) for relatively distantly related taxa for which sequence profiles are available, like mammals and fungi, and using Jackhmmer (52) to map the more recent history of genes among the alveolates.

The three novel CIII proteins (C3AP1-3) have orthologues in *T. gondii*, and one, C3AP1, has an orthologue in *Vitrella brassicaformis*, a sister taxon to the Apicomplexa. They therewith appear to be relatively recent inventions of the Apicomplexa and their close relatives. Three proteins that are present in CIII from fungi and Metazoa and absent from CIII in *P. falciparum, i*.*e*. UQCR1, UQCRC2, and UQCRC10, have all been gained in the evolution of the opisthokonts (53). Only UQCRC11 appears to have been lost specifically from the Apicomplexa (Fig. 4A).

Most of the twelve novel proteins we detected in CIV (Fig. 3B) appear to have a myzozoan origin (Fig. 4b). Of the proteins that are absent from *Pf*CIV, COX5A, COX7B, and COX8 appeared relatively recent in the evolution of CIV in opisthokonts (53). COX7A and COX7C were specifically lost in apicomplexan evolution. These short proteins have a single transmembrane region that, within the 3D structure of CIV in *S. cerevisiae* (54), are in close proximity to each other (Supplementary Fig. 5), suggesting an interlinked alteration on one side of the 3D structure in Apicomplexa. Nevertheless, we cannot exclude that due to sequence divergence they cannot be detected using sequence-based homology detection.

Except for the presence of orthologues in other species, we also examined addition/loss of protein domains in conserved complex members. With respect to CIII, *Pf*CYTC1 contains, relative to CYTC1 in human, an N-terminal extension of ∼150 amino acids that is notable because it is specific to Apicomplexa and is present as an individual protein in *Cryptosporidium muris* (CMU_009920) that lacks a traditional mitochondrion (55). PF3D7_0306500 encodes a 299 amino acid protein of which only the C-terminal ∼50 residues are homologous to COX7A. If, as suggested by homology, PF3D7_0306500 interacts with the other members of CIV in the same manner as COX7A, most of the protein would be located in the matrix. Finally, we detected one novel CIV subunit (ApiCOX13) containing a CCCH zinc finger domain likely harbouring a [2Fe-2S] cluster. ApiCOX13 is orthologous to the human mitochondrial matrix protein CISD3, which plays an important role in iron and ROS homeostasis (56). Despite this direct evolutionary relationship, there are some important differences. CISD3 contains two CCCH zing finger motifs of which only one is conserved in ApiCOX13. Furthermore, ApiCOX13 contains a (predicted) C-terminal transmembrane helix, with a (predicted) topology that puts most of the protein in the mitochondrial matrix, while CISD3 is not a transmembrane protein. When in evolution CISD3 has become part of CIV is not known. The C-terminal transmembrane helix can be detected within the Apicomplexa and in *V. brassicaformis*, but not in the ciliates, suggesting it originated, like many new CIV proteins in *P. falciparum*, in the Myzozoa.

### F_1_F_O_-ATP synthase – Complex V

Classical mitochondrial function includes harnessing energy in the chemical bonds of ATP, a process predominantly executed by CV. In the samples processed with methods 1 and 2 (Supplementary Table 1), we were unable to detect any of the predicted F_1_F_o_-ATP synthase components, including putative apicomplexan-specific subunits recently identified in *T. gondii* (12, 57). The notable exceptions were free-forms of ATPα and ATPβ without any apparent interaction partners. We suspected that this could have been due to harsh lysis conditions, insufficient quantities of mitochondrial protein, inability of the assembled complex to enter the gel or depletion of the complex through saponin treatment (58). To address these issues, larger amounts of ABS parasites or gametocytes were lysed through nitrogen cavitation without saponin (method 3; Supplementary Table 1). To detect protein assemblies >5 MDa, the stacking gel was also analysed. Thus, we found fourteen proteins that are associated with CV, either through homology to previously identified *T. gondii* components or through homology to canonical F_1_ components. These CV subunits comigrated at a size of ∼2.2 MDa or remained stuck at the interface of the stacking gel and the sample slot in both ABS parasites and gametocytes (Fig. 5b). The fraction unable to enter the gel potentially represents higher oligomeric CV states, while we interpret the 2.2 MDa band as an unusually large CV dimer. This is based on a tentative stoichiometry of 10 ATPc subunits for the c-ring, canonical stoichiometry for orthologues to known components and single copies of novel components, predicting a monomer mass of ∼1 MDa (Table 1). This also suggests that the ∼1 MDa complex observed in other studies probably represent the monomer rather than the dimer (7, 11, 59). As the highest abundances were observed in the stacking gel or at the respective monomeric subunit sizes, we plotted iBAQ values for all identified components in the 1-3 MDa range to examine whether intensities in the suspected CV dimer band was comparable. In order to better visualize this, relative abundances were renormalized based on highest iBAQ values in the 1-3 MDa size range (Fig. 5c). We observed similar iBAQ values for the putative CV components around the 2.2 MDa peak, except for ATPα and ATPβ that exhibited about three times higher values consistent with their expected stoichiometries. Furthermore, the abundance peak of ATPdlike was at a lower M_r_ ^app.^ in gametocytes, though the subunit clearly comigrated with the peak of dimeric CV in the ABS parasite sample (Supplementary Fig. 6).

**Figure 5.**
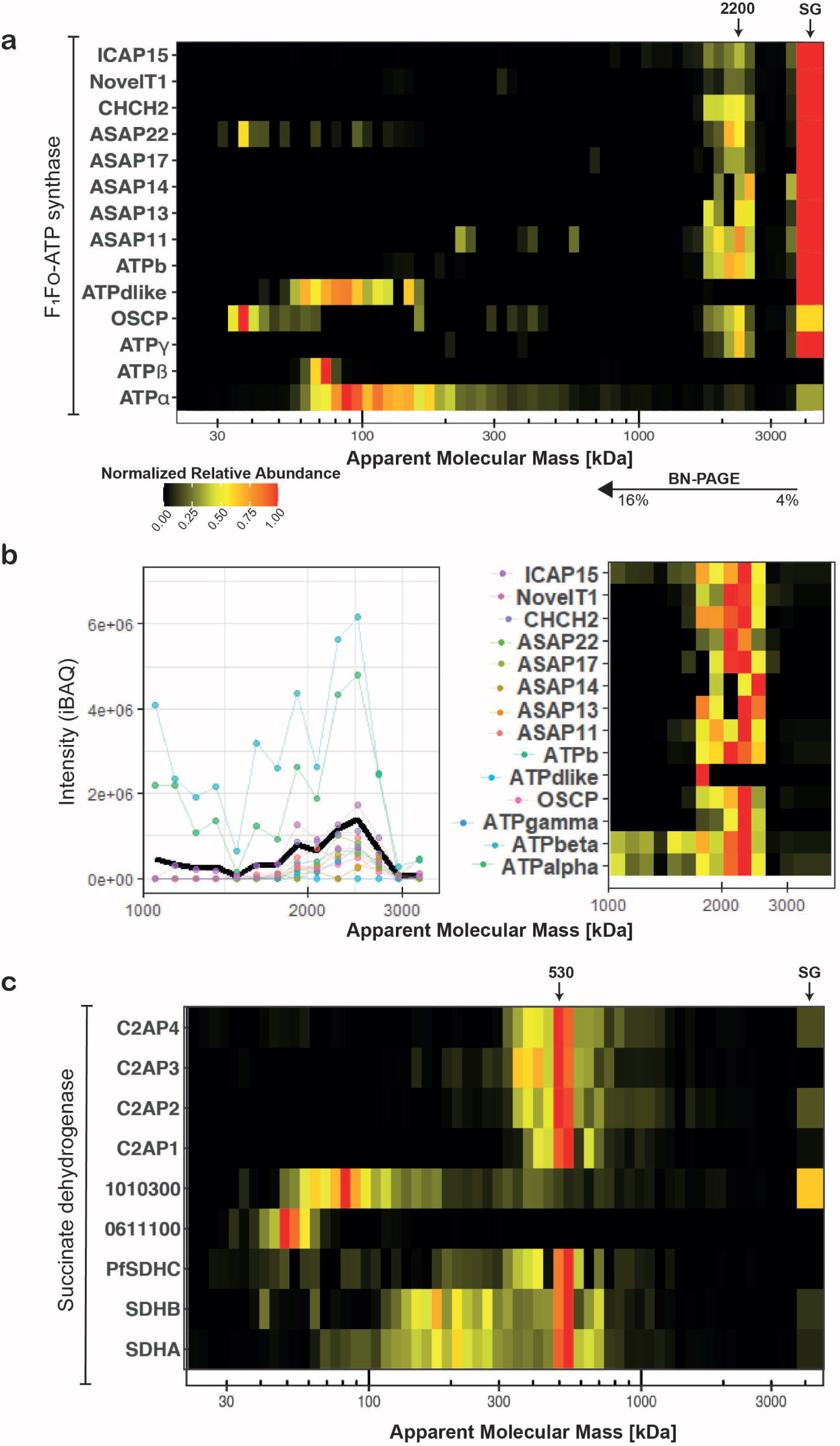
Composition and apparent molecular mass of succinate dehydrogenase (CII) and ATP synthase (CV) in *Plasmodium falciparum* gametocytes. (**a**) Heat map showing migration patterns of canonical ATP synthase components as well as components identified in *T. gondii (12)* in sample GCT3D (67). Most components show comigration at a size band from 2 – 3 Mda as well as abundance in the stacking gel interface (rightmost slice). ATPdlike, ATPβ, OSCP and ATPα appear to be most abundant at their respective monomeric sizes. (**b**) iBAQ values for putative ATP synthase components in 750 – 3500 kDa range (left panel) and heatmap of renormalized ATP synthase data based on intensities for migration at masses >750 kDa, excluding stacking gel interface intensities (right panel).(**c**) Heat map showing SDHA and SDHA comigrating at an M_r_^app.^ of ∼530 kDa along with a group of putative novel components in sample. Stacking gel (SG) is represented broader in the heat map and indicated with a black arrow.

### Succinate dehydrogenase – respiratory chain complex II

Succinate dehydrogenase couples succinate oxidation as part of the citric acid cycle to the reduction of ubiquinone in the OXPHOS pathway. CII is generally composed of at least four different subunits: the hydrophilic SDHA and SDHB subunits catalysing succinate oxidation and the hydrophobic SDHC and SDHD subunits anchoring the complex in the inner mitochondrial membrane and providing the binding pocket for haem and ubiquinone. Similar to CV, we were unable to find an assembled CII using methods 1 or 2. In *Plasmodium*, only SDHA and SDHB are experimentally verified (60). Using method 3, we found comigration of SDHA and SDHB at an M_r_^app.^ of ∼530 kDa (Fig. 5a). Two previously suggested candidates for SDHC (PF3D7_0611100) and SDHD (PF3D7_1010300) (61) were not comigrating with this complex in any of the samples (Fig. 5a). Instead, we identified five putative subunits sharing a common dominant band, although the individual migration patterns were quite heterogeneous and spread over multiple slices. We assigned one candidate as a putative *Pf*SDHC (PF3D7_1448900) as it contains a “DY” motif at positions 52-53 that is conserved in SDHC in a large number of species (62) and of which the tyrosine binds ubiquinone in yeast (63) (Supplementary Fig. 7). However, it contains no recognizable haem-binding motif and only a single (predicted) transmembrane helix, in contrast to three in *S. cerevisiae* SDHC. The other components we named respiratory chain complex 2 associated proteins 1-4 (C2AP1-4; Table 1), one of which (PF3D7_0808450) is myzozoan-specific and has been shown to play a critical role in ookinete mitochondria in *P. berghei* (64). Under native conditions, CII can be found as a trimer in prokaryotes (65, 66). The subunit composition suggested in this study predicts a molecular mass of 188 kDa per CII monomer and 564 kDa for the trimer (Table 1) approximating the observed apparent mass of 530 kDa. As detection of CII components was limited to ABS3D and GCT3D (Supplementary Table 1), sample size is small compared to other complexes discussed in this study. Therefore in this case, further studies will be needed to verify our findings.

### Protein dynamics are in line with a significant metabolic shift in *P. falciparum* gametocytes

Metabolomics approaches have indicated a shift in carbon metabolism in gametocytes from anaerobic glycolysis towards increased TCA cycle utilization and presumably increased respiration (71, 72), which is also reflected in a general increase of mitochondrial proteins and specifically of TCA proteins in gametocytes (73). Likewise, this is supported by an increased sensitivity of gametocytes to TCA cycle inhibition (5) and reliance on wild-type cytochrome *b* for transmission (6), which both are non-essential during ABS. A further indication is the *de novo* appearance of cristae in gametocytes, as they typically serve as hubs for respiration (74) Fig. 1). We examined whether this mitochondrial phenotype would also be reflected in the abundance of OXPHOS complexes.

When normalizing the complexome profiles for total intensity of protein detected in a given fraction, a general trend was observed that proteins associated with CIII and CIV were more abundant in gametocytes than ABS parasites by a large margin (Fig. 3, Supplementary Table 2). However, among different gametocyte samples, abundances of OXPHOS associated proteins also varied up to two-fold even after correction. This indicated that the degree of mitochondrial enrichment was not entirely consistent across the different samples, necessitating a more unbiased approach using stage V gametocyte and mixed ABS parasite whole-cell lysate under denaturing conditions. We performed MS analysis on 30 slices of SDS-gel separated lysate, minimizing false positive identifications by only evaluating the slices that match the predicted molecular mass of the protein. The obtained results complemented and supported essentially all observations made using complexome profiling (Fig. 6). Compared to ABS parasites, the average abundance levels of OXPHOS complex components in gametocytes were higher 15-fold for CII, 36-fold for CIII, 44-fold for CIV, and 32-fold for CV. This phenomenon included all but one of the putative novel components, further supporting their association with the respective complexes. Outliers were PFSDHN3 from CII, CYTB from CIII, and COX2a from CIV, which showed a comparatively higher abundance in ABS parasites. For subunit COX2a, one identified peptide with a high error and inconsistent migration pattern was observed. After manual removal of this peptide and recalculation, the iBAQ value was in line with the other CIV components. For the other two proteins no obvious outliers were observed at the peptide level. Therefore, this data does not support CII membership of C2AP3. It is noteworthy that all outliers were proteins detected with a low peptide count and few MS/MS events, suggesting decreased reliability when attempting to quantify proteins close to the detection limit.

**Figure 6.**
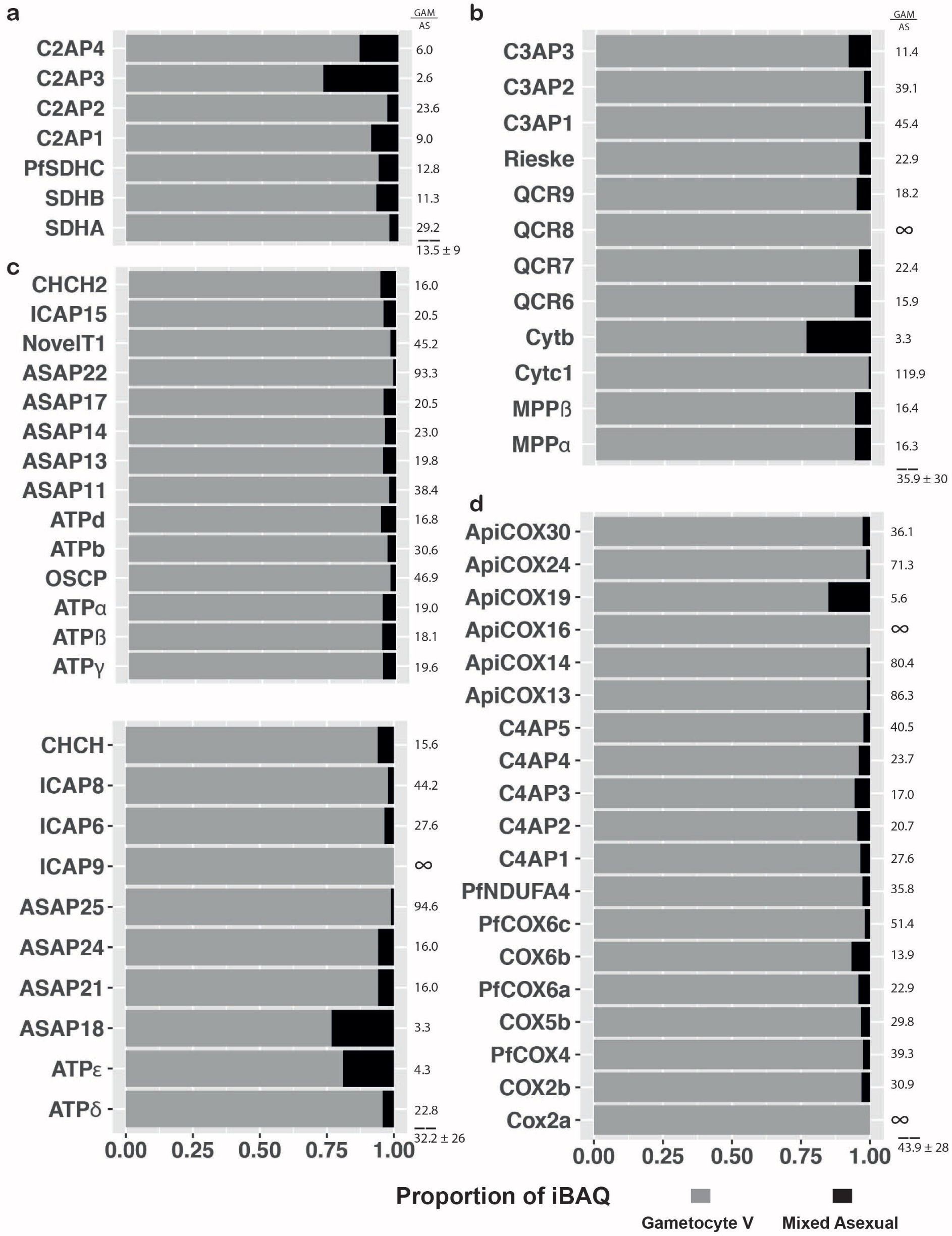
Relative quantification of respiratory chain complex components. Relative abundance expressed in proportion of iBAQ of OXPHOS components in ABS parasites (black) and gametocytes (grey). Data is based on denatured whole cell lysates separated by SDS-PAGE and analysed by label-free quantitative MS. Fold changes are indicated next to each bar and averages with standard deviation for components of each complex are indicated below dashed lines. Infinite fold changes were arbitrarily treated as 100 for average/SD calculations. (**a**) Putative components of CII. (**b**) Putative components of CIII. (**c**) Putative components of ATP synthase detected in complexome profiles (upper section) or only detected in SDS profiles but identified as an ATP synthase component in *T. gondii*(12, 67) (lower section). (**d**) Putative components of CIV.

To further analyse whether this trend was indicative of a larger metabolic shift, we also investigated abundance dynamics of other proteins involved in central energy metabolism (Fig. 7). Complexome (Supplementary Fig. 8) and SDS profiles (Fig. 7a) indicated that enzymes involved in the glycolysis pathway are much more prevalent in ABS parasites. Interestingly, an alternative lactate dehydrogenase (altLDH; PF3D7_1325200) – or potentially malate dehydrogenase as substrate specificity cannot be deduced from sequence – appears to be gametocyte-specific. This is also true for an alternative phosphofructokinase (altPFK; PF3D7_1128300) that appears to have lost crucial residues required for its function (75). Stage differences varied for enzymes of the TCA cycle, potentially suggesting different inputs, bottlenecks or alternative utilization of individual enzymes between stages (Fig. 7b), however, these did not correlate with gene essentiality (5). Alternative ubiquinol producing enzymes that feed into OXPHOS were also increased in gametocytes but to a lesser degree for DHODH, which had a comparatively higher abundance in ABS parasites (Fig. 7d). DHODH is expected to be abundant in ABS parasites due to their reliance on de novo pyrimidine biosynthesis (4). All mitochondrial enzymes were found to be comparatively more prevalent in ABS parasites when assessed through mitochondria-enriched complexome profile data (Supplementary Fig 8). This is possibly due to higher relative mitochondrial content in those samples as is suggested by the comparatively higher abundance of mitochondrial “household” genes VDAC, TOM40, TIM50 and HSP60 (Fig. 7c). Taken together these data support previous metabolomics-based suggestions of a switch towards respiration and away from anaerobic glycolysis in *P. falciparum* gametocytes (71).

**Figure 7.**
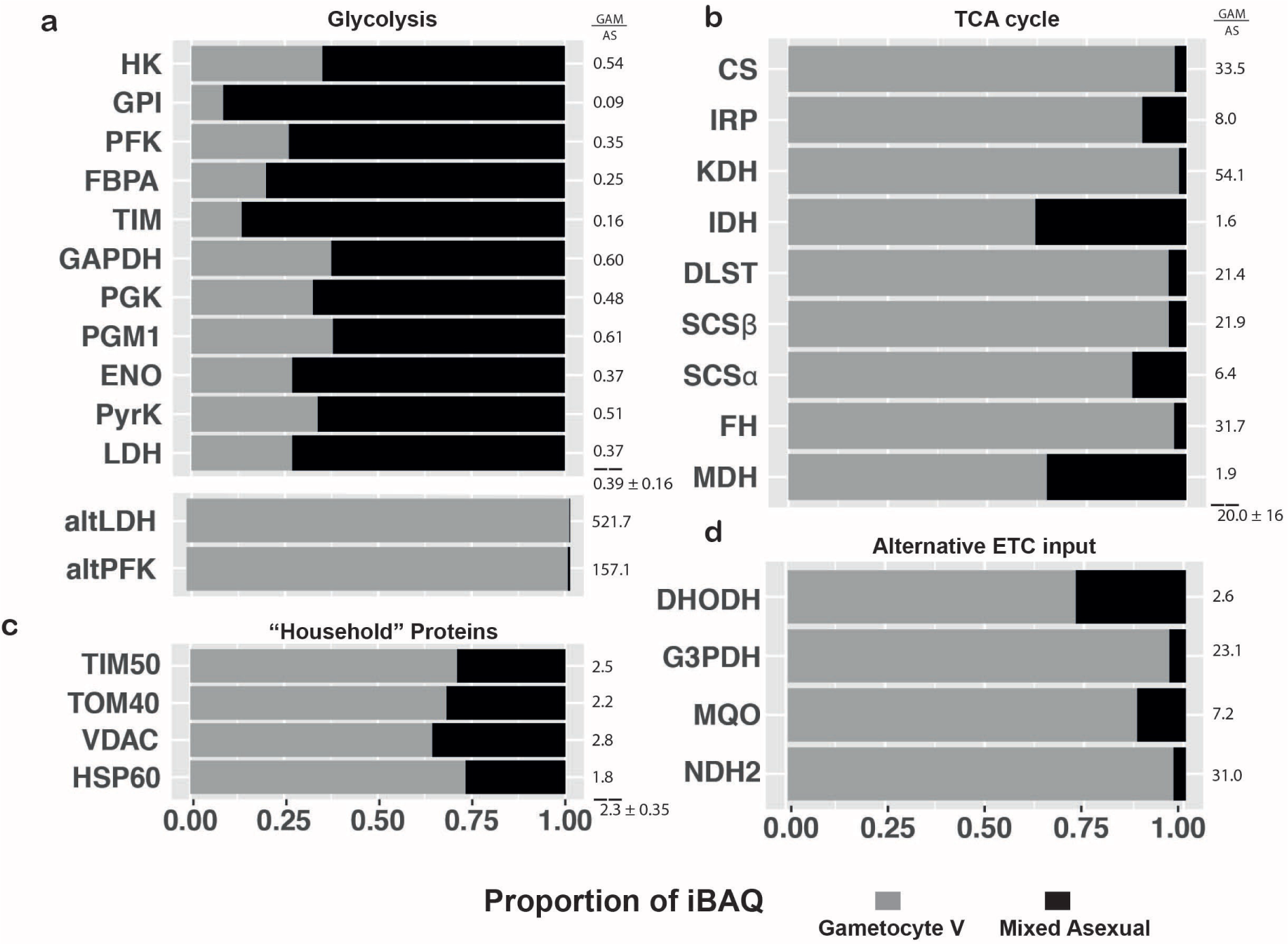
Abundance comparison of energy metabolism related proteins. Relative quantification of a selection of energy metabolism enzymes (**a, b, d**) and mitochondrial household proteins (**c**) in ABS parasites (black) and gametocytes (grey) based on denatured whole cell lysates separated by SDS-PAGE. Fold changes are indicated next to each bar and averages with standard deviation for a group are indicated below dashed lines. Corresponding Gene IDs can be found in Supplementary Information S1

## Discussion

We demonstrated the utility of complexome profiling to address the considerable knowledge gap regarding multiprotein assemblies and supercomplex formation in *P. falciparum*. We identified putative novel OXPHOS complex components, suggested a mechanism for regulation of crystalloid formation in *P. falciparum*, and validated and predicted additional features for previously characterized complexes. Additionally, we utilized the label-free quantification data from complexome profiling and denatured whole cell lysates to assess abundance changes between ABS and mature gametocytes. To place our data into an evolutionary context, we integrated phylogenetic analysis allowing us to devise novel hypotheses and assess significance of observed phenomena.

For the novel uncharacterized and apicomplexan-specific OXPHOS complex subunits, it is challenging to estimate biological significance or function. It is tempting to speculate that some of the new, transmembrane helix containing proteins are functional and structural replacements of the two missing subunits in CIV, however, as we have observed in the evolution of CI in Metazoa, evolutionary new subunits do not necessarily occupy the same location in the complex where subunits are missing (76). Nevertheless, the observed complexes allow us to draw some important conclusions and raise interesting questions. CIII provides a clear example where *Plasmodium spp*. resemble plants by both using MPPα and β as structural components (48) unlike animals and fungi where MPPα and β have been replaced by the homologous subunits core 1 and 2 that do not have general MPP activity. The fact that the mitochondrial processing peptidases are tied to the structurally essential core 1 and 2 subunits of cytochrome bc_1_ complex, presents an interesting trade-off in the context of *Plasmodium* biology. *P. falciparum* ABS parasites are not reliant on OXPHOS outside of ubiquinone recycling for pyrimidine biosynthesis (4), while gametocytes heavily rely on it for successful colonization of and development in the insect host (6, 7). As a direct consequence, we see a much lower specific content of OXPHOS complexes in ABS parasites compared to gametocytes (Fig. 6). However, presumably ABS parasites still have a comparably high need for MPP activity to facilitate mitochondrial function outside of respiration, as exemplified by putative MPPβ inhibitors showing promise as antimalarials (77, 78). This would necessitate synthesis of the whole complex, which is much less efficient and more challenging from a regulatory standpoint than generating the heterodimer observed in the mitochondria of its host. A dual localization of the processing peptidases as a soluble heterodimer would circumvent this but is not observed under our conditions.

Similarly puzzling, there is an apparent increase in size and number of components of OXPHOS complexes, despite the minimalistic mitochondrial genome and single subunit type II NADH:ubiquinone oxidoreductase replacing CI. In most commonly studied eukaryotes, a typical M_r_^app.^ is ∼500 kDa for CIII dimer, ∼210 kDa for CIV monomer, ∼120 kDa for CII monomer, and ∼600 kDa for CV monomer. Our data suggest respective size increases of around 50%, 130%, 50% and 70%, as compared to complexes III, IV, II and V of more commonly studied eukaryotes. While enlarged OXPHOS complexes have been observed before (53), the size increases in *P. falciparum* are remarkably large. Furthermore, these increases occurred relatively late in evolution, either around 950 million years ago, before the origin of the Myzozoa, or 830 million years ago, before the origin of the Apicomplexa (79). In contrast, fungi and Metazoa have comparably few of such taxon-specific supernumerary subunits. Without further experimental investigation, it is challenging to say whether these size increases correspond to additional functions of the OXPHOS complexes such as the earlier discussed MPP activity. An alternative explanation could be that the transfer of genes originally found on mtDNA to the nucleus necessitated amino acid changes to facilitate import into the mitochondrion, which in turn required incorporation of additional subunits to maintain the same functionality. Although this is a tempting hypothesis, no strong evidence for it has been obtained from the analysis of the evolution of the mitochondrial genome in conjunction with the mitochondrial proteome in other evolutionary lineages (76, 80). Finally, there also could be a lack of energetic constraints that are afforded by the parasitic lifestyle, which may have allowed the passing on of such bulky complexes. A comparable phenomenon was shown in lower mitochondrial quality control and selective constraints of flightless birds compared to their flying counterparts that retained this energetically demanding ability (81).

In the context of *de novo* cristae biogenesis in gametocytes, possible consequences of changes in the abundance of CV dimers and their markedly different composition are particularly notable: rows of CV dimers (82) are known to shape cristae by bending their membrane (83). The observed decrease to ∼3% of CV components in ABS compared to gametocytes (Fig. 6) thus provides a straightforward explanation for the absence of cristae in the former. The large amounts of ATP synthase found in the stacking gel interface (Fig. 5, Supplementary Fig. 6) may suggest a particular stability of higher order CV assemblies. Possibly this is conferred by the additional subunits found in *P. falciparum*, which may also be responsible for the unusual shape of the cristae in the parasite.

In summary, our comprehensive comparative analysis of the ABS and gametocyte complexome profiles has revealed abundant clade-specific novelties and overwhelming stage-differences. A further fundamental understanding of these differences could help to leverage the heavy reliance of gametocytes on unusual and highly divergent mitochondrial complexes as much sought-after gametocytocidal drug targets. To this end, application of genetic tools will be a crucial next step to assess the role and significance of the divergent features proposed in this study. Above all, complexome profiling of *P. falciparum* mitochondria revealed peculiar new biology and fascinating insights in the evolution of eukaryotic respiration and how the malaria parasite has adapted to different environmental challenges at the level of multiprotein complexes.

## Material and Methods

### Parasite culture

*P. falciparum* strain NF54 was maintained in RPMI [7.4] supplemented with 10% human serum and 5% haematocrit using standard culturing technique as described previously (84). NF54/iGP2 strain was additionally supplemented with 2.5 mM D-(+)-glucosamine hydrochloride (Sigma #1514) for maintenance. For induction, parasites were synchronized with 5% sorbitol as described previously (85) and glucosamine was omitted from the medium for 48 hours. From day 4-8 gametocyte cultures were treated with 50 mM N-acetylglucosamine to eliminate ABS parasites as described previously (86). For WT-NF54 gametocytes induction the same procedure was followed except that instead of glucosamine induction, parasites were overgrown for 5 days with only medium exchanges every 48 hours prior to N-acetylglucosamine treatment. At day 14 after induction, gametocytes were magnet-purified according to the procedure described previously (87). ABS parasites were not further enriched prior to processing. Infected red blood cells were freed from host material through 10 min incubation in 10x pellet volume of 0.05% (w/v) saponin in phosphate buffered saline (PBS; pH7.4) and subsequent centrifugation at 3000 x g for 5 min. The parasite pellet was washed twice with PBS and the dry pellet was flash frozen and stored at −80°C.

### Transmission electron microscopy

For electron microscopy analysis of mitochondria across different asexual blood-stage parasites and mature gametocytes, infected red blood cells were fixed in 2% glutaraldehyde in 0.1 M cacodylate (pH 7.4) buffer overnight at 4°C, washed and cell-pellet was resuspended in 3% ultra-low-gelling agarose, solidified and cut into small blocks. Agarose blocks with cells were postfixed for 1 h at RT in 2% osmium tetroxide and 1.5% potassium ferrocyanide in 0.1 M cacodylate buffer with 2mM CaCl2, washed in MQ and incubated in 0.5% thiocarbohydrazide solution for 30 min at RT. After washing agarose blocks with cells were again fixed in 2% osmium for 30 min at RT, washed and placed in 2% aqueous uranyl acetate overnight at 4°C. After washing agarose blocks with cells were placed in lead aspartate solution (pH 5.5) for 30 min at 60°C, washed, dehydrated in an ascending series of aqueous ethanol solutions and subsequently transferred via a mixture of aceton and Durcupan to pure Durcupan (Sigma) as embedding medium. Ultrathin sections (80 nm) were cut, air dried and examined in a JEOL JEM1400 electron microscope (JEOL) operating at 80 kV.

### Mitochondrial enrichment

On the day of the experiment, parasite pellets were resuspended in ice-cold MESH buffer (250 mM sucrose, 10 mM HEPES, 1 mM EDTA, 1x cOmplete™ EDTA-free Protease Inhibitor Cocktail (Sigma), pH 7.4) and washed by centrifugation at 3500 x *g*; 10 min, 4 °C. The mitochondria-enriched fractions were obtained following three different methods. For *methods 1 and 2*, the parasite pellets were resuspended in ice-cold MESH buffer supplemented *with* or *without* 0.5% (w/v) saponin, respectively and lysed by 20 strokes through a 27G needle. Rough debris and unbroken cells were pelleted at low speed centrifugation (600 x *g*, 10 min, 4 °C). The supernatant was transferred into a new tube and a low speed centrifugation was repeated. The supernatant was recovered again and centrifuged at a higher speed (22000 x *g*, 15 min, 4 °C). The supernatant (cytosolic fraction) was discarded and the pellet (mitochondria-enriched fraction) was resuspended in MESH buffer and kept on ice until usage. For *method 3*, nitrogen cavitation was used for cell disruption. The parasite pellets were washed once in ice-cold MESH, pooled in a total volume of 1 ml and then added to a pre-chilled cell disruption vessel (#4639 Parr Instrument Company). The vessel was pressurized and equilibrated with nitrogen gas at 1500 psi for 10 minutes on ice. The parasite cells were then sheared through a slow release by nitrogen cavitation. The mitochondria-enriched fraction was obtained by differential centrifugation as described above. Protein concentration was determined using the Pierce™ BCA Protein Assay Kit (Thermo Scientific) using bovine serum albumin as standard.

### Blue Native Polyacrylamide Gel Electrophoresis (BN-PAGE)

Protein samples (∼150 μg) were resuspended in 500 mM 6-aminohexanoic acid, 1 mM EDTA and 50 mM imidazole/HCl (pH 7.0) and solubilized with either Triton X-100 (Sigma), Digitonin (SERVA) or n-dodecyl-β-D-maltoside (DDM) (Sigma) using detergent:protein (w/w) ratios of 10:1, 6:1 and 3:1, respectively. The solubilized samples were centrifuged at 22000 x *g* for 20 minutes; 4 °C. The supernatants were recovered, supplemented with Coomassie-blue loading buffer and separated on 4%–16% polyacrylamide gradient blue native gels as described previously(18). For mass calibration, purified bovine heart mitochondria (50-100 μg) solubilized under the same conditions were run alongside each set of *Plasmodium* samples.

### Denaturing polyacrylamide gel electrophoresis (SDS-PAGE)

Mature gametocyte or ABS parasite pellets were generated according to the procedure described above. Samples were lysed in SDS loading buffer(1% β-Mercaptoethanol, 0.004% Bromophenol blue, 6% glycerol, 2% SDS 50mM Tris-HCl, pH 6.8) and heated for 5 minutes at 95°C. Insoluble debris were pelleted by centrifugation (20000 x G, 5min, RT). Supernatants were recovered and 20 μg protein were separated on SurePAGE Bis-Tris 4-12% gradient gel following the manufacturer’s instructions (Genscript). A protein standards ladder (BioRad #1610374) was run alongside the samples for mass calibration.

### In-Gel Trypsin Digestion

After electrophoresis, the gels were fixed in 50% methanol, 10% acetic acid, 10 mM ammonium acetate, stained with 0.025% Coomassie blue G-250 (SERVA) in 10% acetic acid for 30 min, destained in 10% acetic acid and kept in deionized water. A real size colour picture was taken using an ImageScanner III (GE Healthcare) to prepare a template for the cutting procedure. The in-gel tryptic digestion was carried out following the method described in Heide *et al*. 2012(88) with slight modifications. In brief, each gel lane was cut into 30 or 60 even slices starting at the bottom. Each slice was further diced into smaller pieces before being transferred to a filter microplate (96 wells, Millipore MABVN1250) prefilled with 200 μl 50% methanol, 50 mM ammonium hydrogen carbonate (AHC) per well. In order to remove the Coomassie dye, the gel pieces were incubated in the same solution at room temperature (RT) and washed by centrifugation (1000 x *g*, short spin) until flow through was clear. For cysteines reduction, the gel pieces were incubated in 10mM DL-dithiothreitol (DTT), 50mM AHC for 60 minutes at RT under gentle agitation. The solution was removed by centrifugation at 1,000 x *g*; short spin. In the next step, for cysteines alkylation, the gel pieces were incubated in 30 mM 2-chloroacetamide (CAA), 50 mM AHC for 45 min at RT and solution was removed as above described. The gel pieces were dehydrated in 50% methanol, 50 mM AHC for 15 min. Solution was removed and gel pieces were dried at RT for 45 min at RT. Then, 20 μl of 5 ng/μl sequencing grade trypsin (Promega), 50mM AHC, 1mM CaCl_2_ were added to the dried gel pieces, incubated at 4°C for 30 minutes before 50 μl mM AHC were added and incubated overnight at 37°C in a sealed bag. The next day, the peptides were collected in 96-well PCR microplates by centrifugation at 1000 x *g*; 15 s. The remaining peptides were eluted by incubating gel pieces with 50 μl 30% acetonitrile (ACN), 3% formic acid (FA) for 15 minutes at RT under gentle agitation and collected in the same PCR microplates. The peptide-containing solution was dried in a SpeedVac Concentrator Plus (Eppendorf) and the dried peptides were resuspended in 20 μl 5% ACN, 0.5% FA and stored at −20°C for subsequent analysis.

### Mass Spectrometry

Resulting peptides were separated by liquid chromatography (LC) and analysed by tandem mass spectrometry (MS/MS) in a Q-Exactive mass spectrometer equipped with an Easy nLC1000 nano-flow ultra-high-pressure liquid chromatography system (Thermo Fisher Scientific). Briefly, peptides were separated using a 100 μm ID × 15 cm length PicoTip emitter column (New Objective) filled with ReproSil-Pur C18-AQ reverse-phase beads of 3 μm particle size and 120 Å pore size (Dr. Maisch GmbH) using linear gradients of 5%–35% acetonitrile, 0.1% formic acid (30 min), followed by 35%-80% ACN, 0.1% FA (5 min) at a flow rate of 300 nl/min and a final column wash with 80% ACN (5 min) at 600 nl/min. The mass spectrometer was operated in positive mode switching automatically between MS and data-dependent MS/MS of the top 20 most abundant precursor ions. Full-scan MS mode (400– 1,400 m/z) was set at a resolution of 70,000 m/Δm with an automatic gain control target of 1 × 10^6^ ions and a maximum injection time of 20 ms. Selected ions for MS/MS were analysed using the following parameters: resolution 17,500 m/Δm, automatic gain control target 1 × 10^5^; maximum injection time 50 ms; precursor isolation window 4.0 Th. Only precursor ions of charge z = 2 and z = 3 were selected for collision-induced dissociation. Normalized collision energy was set to 30% at a dynamic exclusion window of 60 s. A lock mass ion (m/z = 445.12) was used for internal calibration(89).

### Complexome profiling

Raw MS data files from all slices were analysed using MaxQuant (v1.5.0.25)(90). For protein group identification peptide spectra were searched against a *P. falciparum* reference proteome (isolate 3D7, version March 21, 2020) as well as a list of common contaminants; e.g. BSA and human keratins. Standard parameters were set for the searches, except for the following: N-term acetylation and methionine oxidation were allowed as variable modifications; up to two missed trypsin cleavages were allowed; cysteine carbamidomethylation as fixed modification; matching between runs was allowed and 2 min as matching time window; FDR as determined by target-decoy approach was set to 1%; 6 residues as minimal peptide length. To allow for abundance comparisons between samples, label-free quantification was applied to each detected protein in the form of intensity based absolute quantification (iBAQ) values. Potential differences in protein quantity and instrument sensitivity between runs were corrected by normalizing for the sum of total iBAQ values from each sample. Protein migration profiles were hierarchically clustered by an average linkage algorithm with centred Pearson correlation distance measures using Cluster 3.0(91). Further analysis of the complexome profiles consisting of a list of proteins arranged depending on their similar migration patterns in the BN gel was performed in R(92) and the results were visualized using ggplot2(93). The mass calibration was performed using the known masses of the mitochondrial oxidative phosphorylation complexes in bovine heart: CII (123 kDa); CIV (215 kDa); CIII (485 kDa); CV (700 kDa); CI (1000 kDa); supercomplex I-III (S_0_, 1500 kDa); supercomplex I-III-IV (S_1_, 1700 kDa); supercomplex I-III-IV_2_ (S_2_, 1900 kDa)

### Homology detection

In order to detect homologous proteins, we used profile-based sequence analysis tools. The sequence profile of each protein was queried between human, yeast, *Arabidopsis, Toxoplasma* and *Plasmodium* proteomes and back, with its respective best hit, using HHpred (51). Orthology was thus confirmed when retrieving the original query. Independently, these proteins were utilized to perform profile-based sequence analysis against the UniProtKB with JACKHMMER(94) in order to find orthologs among the species listed, examining always consistencies with aforementioned findings.

## Acknowledgements

We thank the molecular team members of the Malaria Research Group for fruitful discussions. We also thank Sergio Guerrero-Castillo for his support and helpful discussions. Furthermore, we would like to thank Yoeri van Strien for helping in data exploration, helpful discussions and setting up CEDAR. F.E. and T.W.A.K. are supported by the Netherlands Organisation for Scientific Research (NWO-VIDI 864.13.009), A.C.O., U.B. and M.H. by the Netherlands Organization for Health Research and Development (TOP 91217009) and U.B. by the Netherlands Organization for Scientific Research (TOP 714.017.004).

## Author Contributions

F.E. and A.C.O. performed experiments and analysed results. M.K.L. performed transmission electron microscopy. D.M.E. and M.A.H. performed phylogenetic analysis and contributed illustrations. S.D.B and T.S.V provided the NF54/iGP2 line. U.B. provided conceptual advice and resources. F.E. prepared illustrations and wrote the first manuscript draft. T.W.A.K. conceived and designed the study, provided resources and edited the manuscript. All authors contributed to data interpretation and provided feedback on the manuscript. All authors approved the final version of the manuscript.

## Data Availability

All raw and processed complexome data generated in this study was deposited at the ComplexomE profiling DAta Resource (CEDAR) and can be retrieved under www3.cmbi.umcn.nl/cedar/browse/experiments/CRX23.

## Supplement

**Supplementary Table 1.** Description of all samples analysed by complexome profiling.

| Sample Name | Parasite Stage | Parasite strain | Detergent: Protein | Protein Quantity [μg] | Enrichment method |
| --- | --- | --- | --- | --- | --- |
| GCT1Da | Stage V Gametocyte | NF54 | Digitonin 6:1 | 115 | 1 |
| GCT1Db | Stage V Gametocyte | NF54/iGP2 | Digitonin 6:1 | 100 | 1 |
| GCT1Dc | Stage V Gametocyte | NF54/iGP2 | Digitonin 6:1 | 180 | 1 |
| GCT3D | Stage V Gametocyte | NF54/iGP2 | Digitonin 4.5:1 | 400 | 3 |
| ABS1Da | Mixed ABS | NF54/iGP2 | Digitonin 6:1 | 150 | 1 |
| ABS1Db | Mixed ABS | NF54/iGP2 | Digitonin 6:1 | 150 | 1 |
| ABS2D | Mixed ABS | NF54 | Digitonin 6:1 | 150 | 2 |
| ABS3D | Mixed ABS | NF54/iGP2 | Digitonin 4.5:1 | 300 | 3 |
| ABS1Ma | Mixed ABS | NF54/iGP2 | DDM 3:1 | 150 | 1 |
| ABS1Mb | Mixed ABS | NF54/iGP2 | DDM 3:1 | 150 | 1 |
| ABS2M | Mixed ABS | NF54 | DDM 3:1 | 150 | 2 |

**Supplementary Table 2.**
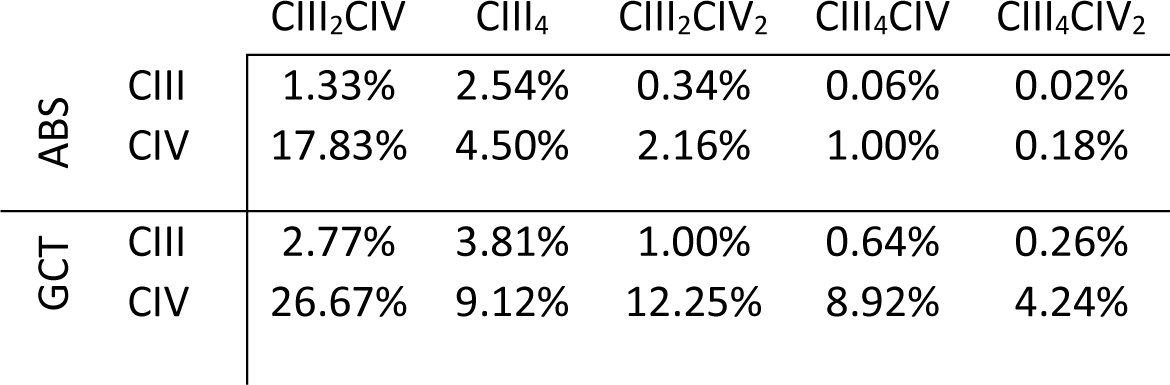
Abundance of putative supercomplex in asexuals and gametocytes normalized separately. Intensities at were normalized against highest value found in ABS3D (ABS) and GCT3D (GCT) respectively and averaged for all complex components. Relative abundance compared to highest intensity value was expressed in %.

**Supplementary Figure 1.**
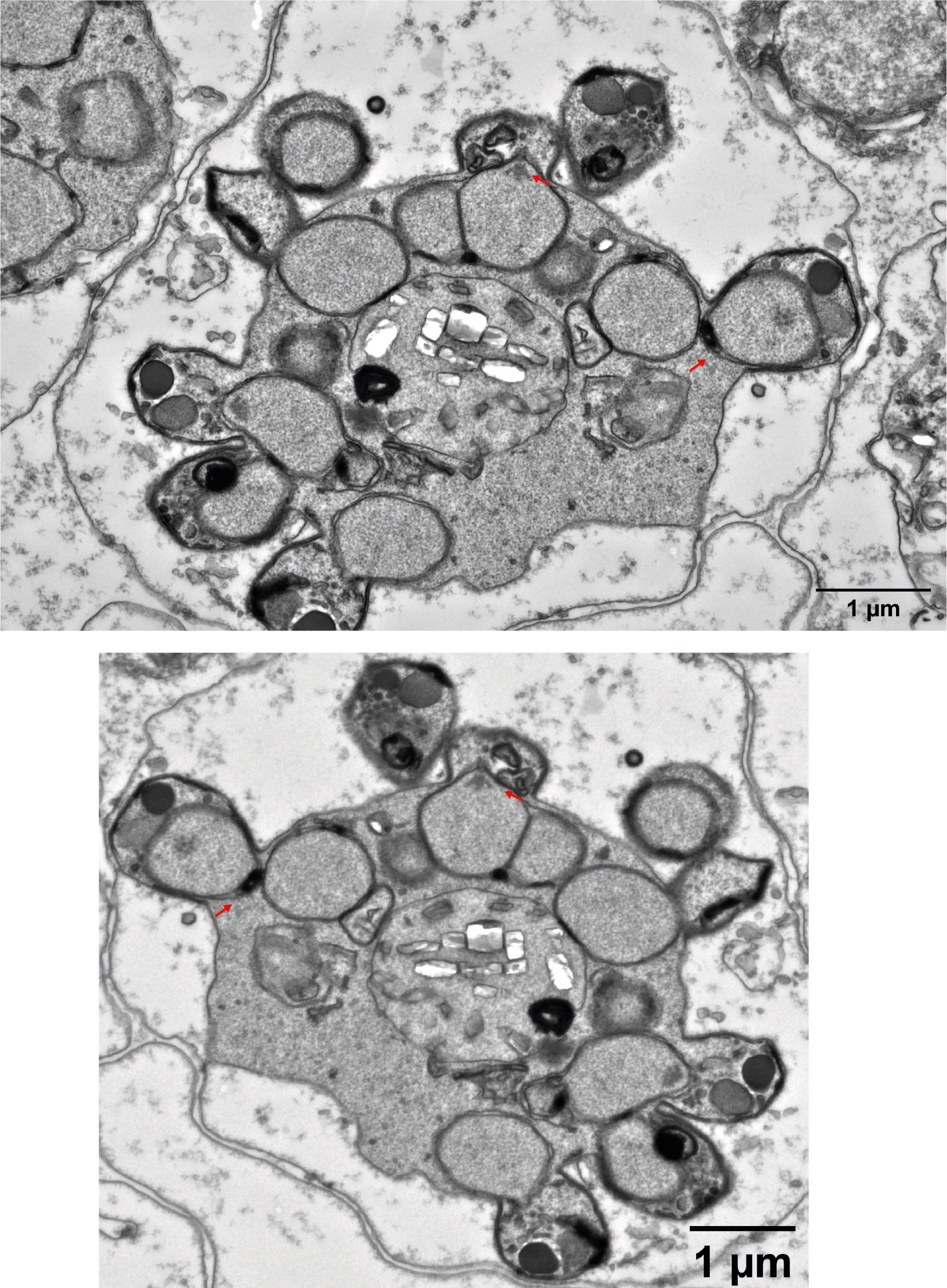
Saponin-lysed infected red blood cells containing early schizonts. Nuclei appear to enter merozoite compartment after organelles, closing off the compartment after entering (red arrows). More electron-dense part of nucleus during entrance process also potentially indicates directed “pulling” of the nucleus.

**Supplementary Figure 2.**
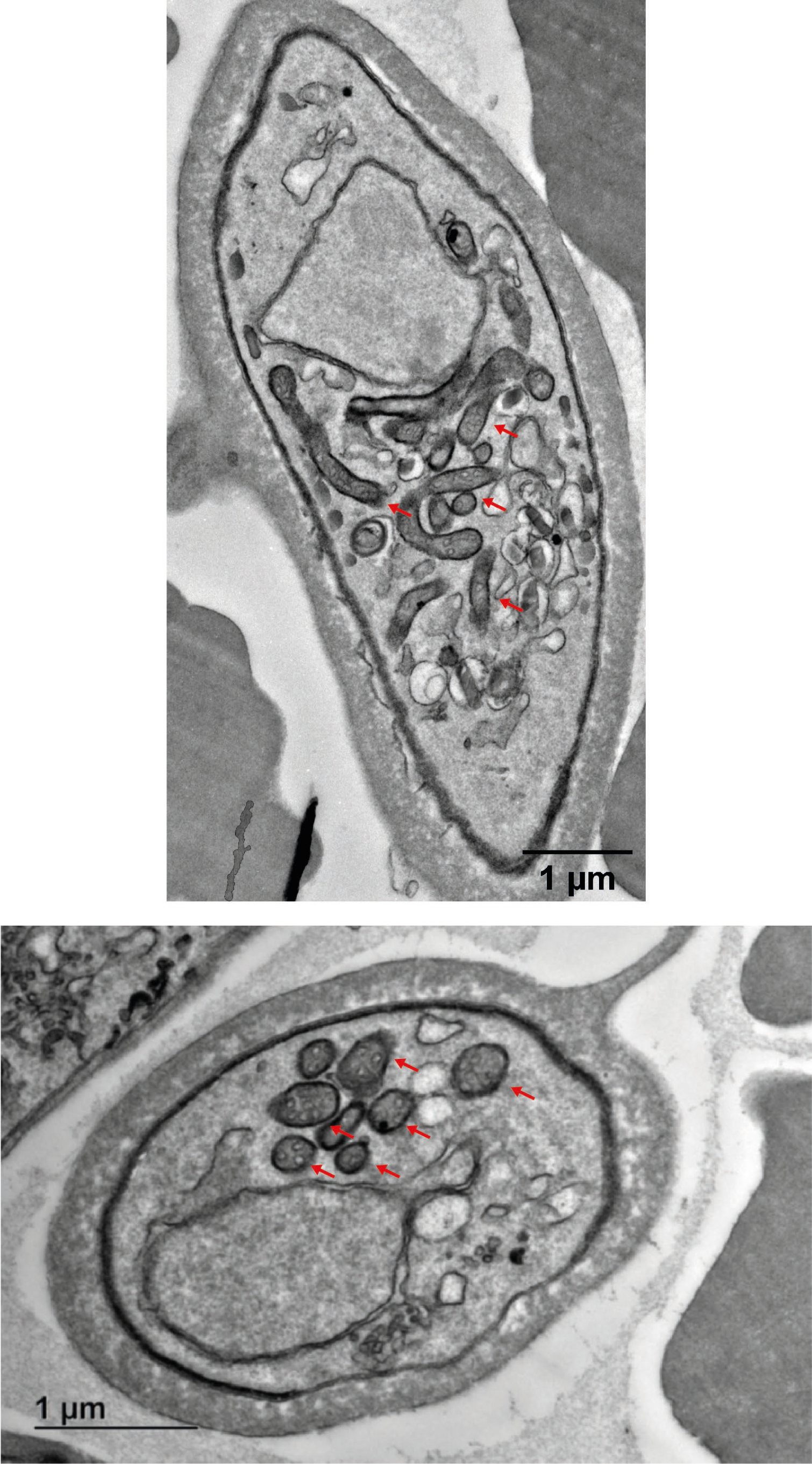
Cristate mitochondrial sections cover large proportion of mature gametocytes. Red arrows indicate mitochondrial sections. Upper panel depicts longitudinal section of a gametocyte and bottom panel depicts horizontal section of a gamteocyte.

**Supplementary Figure 3.**
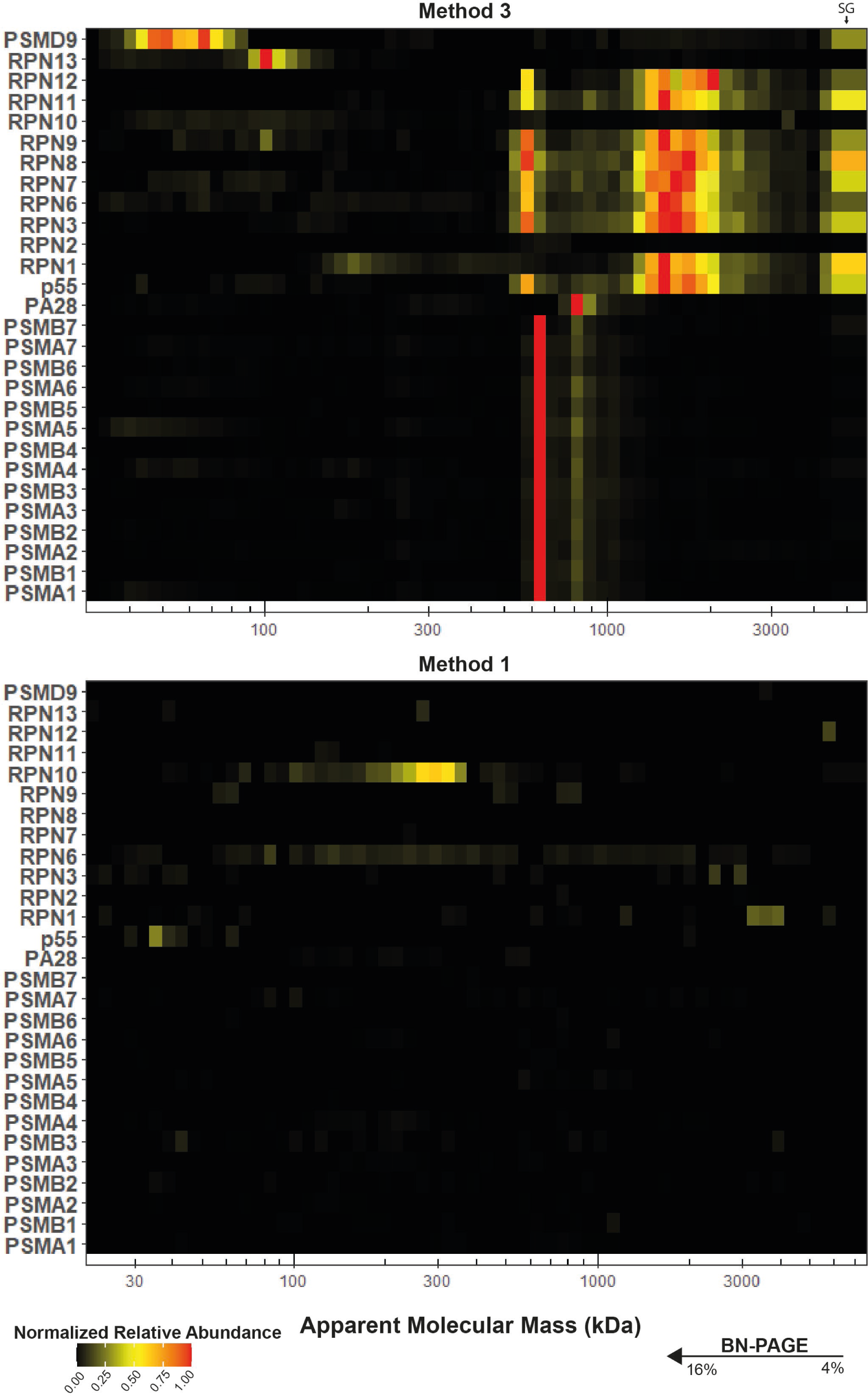
Differential presence of assembled proteasome components. Upper panel depicts abundance of proteasome components in sample ABS3D (method 3), lower panel depicts abundance in sample ABS1Da (method 1). Samples were based on highest iBAQ value for each protein group between the two samples.

**Supplementary Figure 4.**
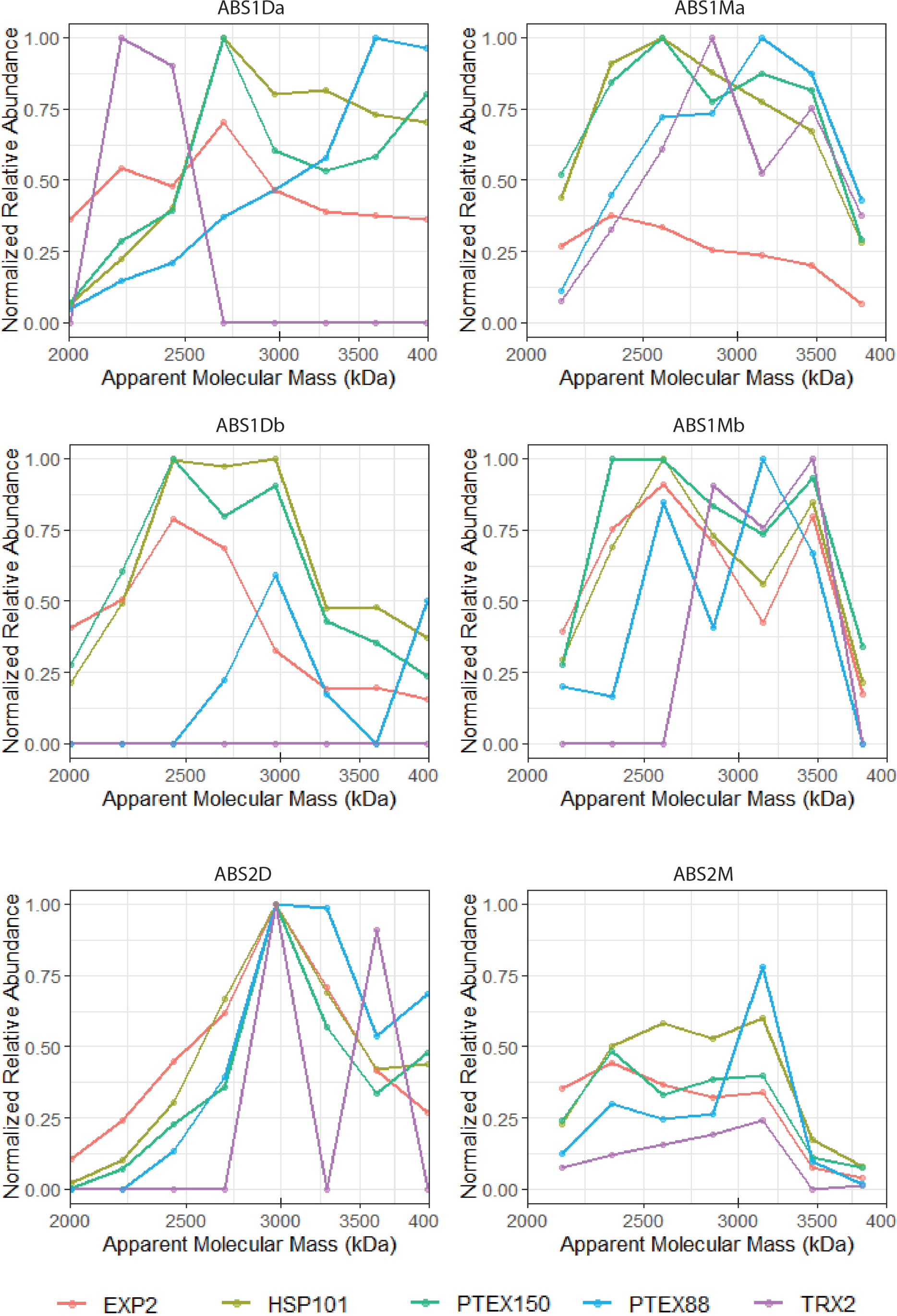
Heterogeneous migration comigration of PTEX88 and TRX2 with PTEX core. Proteins were normalized based on highest iBAQ value in 1500-4000 kDa mass range.

**Supplementary Figure 5.**
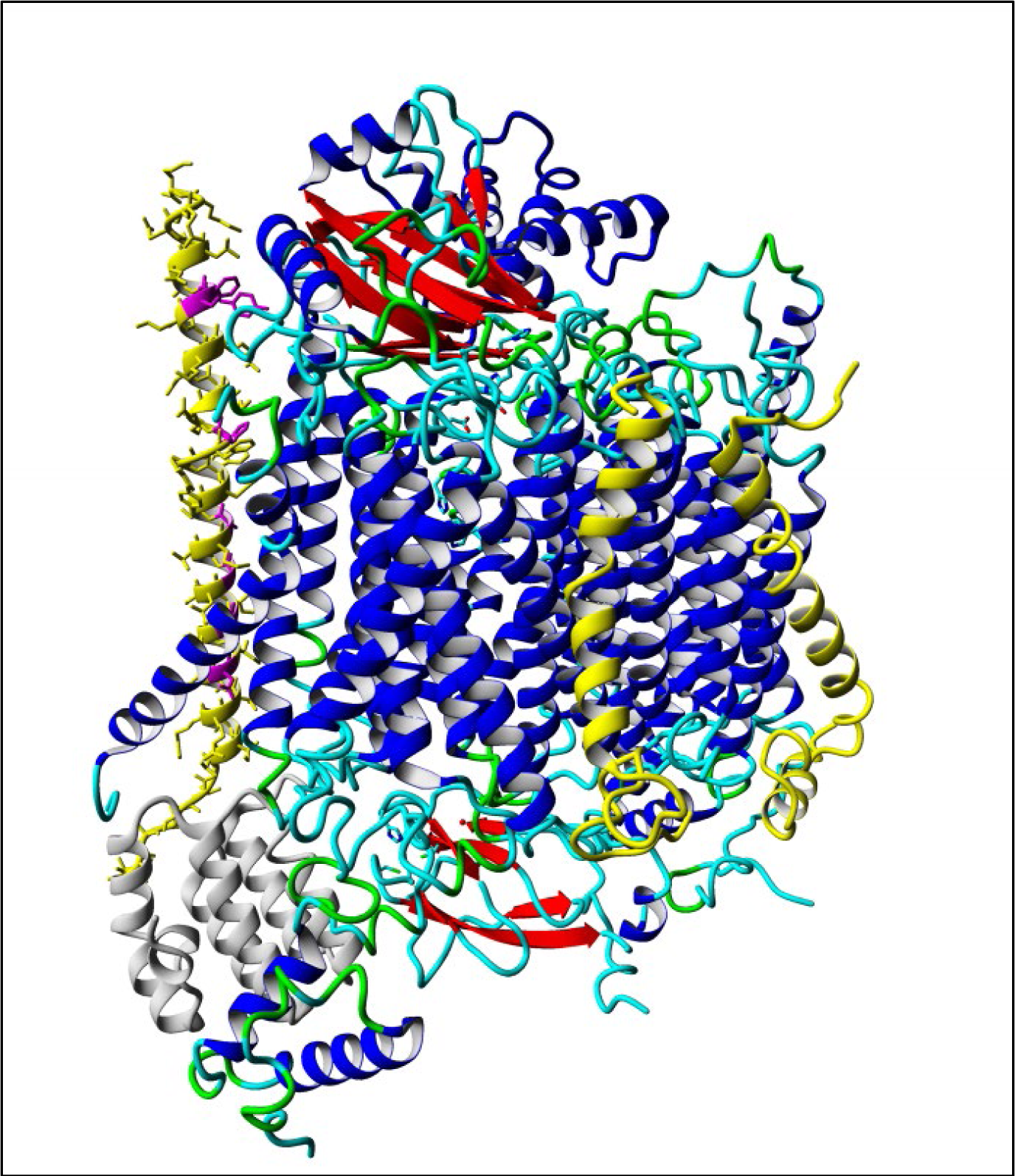
CIV from *S.cerevisiae (95)*. Subunits that were lost in the evolution to *P*.*falciparum*, or whose homology to *P. falciparum* proteins is barely detectable, are in yellow. The gray subunit on the bottom-left (named COX6 in *S*.*cerevisiae* that corresponds to COX5A in metazoa) is present in *S*.*cerevisiae* but is an evolutionary addition that is specific to the opisthokonts and was thus never “lost”. The yellow subunit on the left (COX9 in S. cerevisiae that corresponds to COX6C in metazoa) is poorly conserved in *P. falciparum*. Only the residues in magenta, which appear to interact with other proteins of the complex, are conserved. The subunits that date back to the last eukaryotic common ancestor and for which no homologs could be detected in *P. falciparum* are COX8 (COX7C in metazoa) in the middle and COX7 (COX7A in metazoa) on the right. Visualization done with Yasara (96).

**Supplementary Figure 6.**
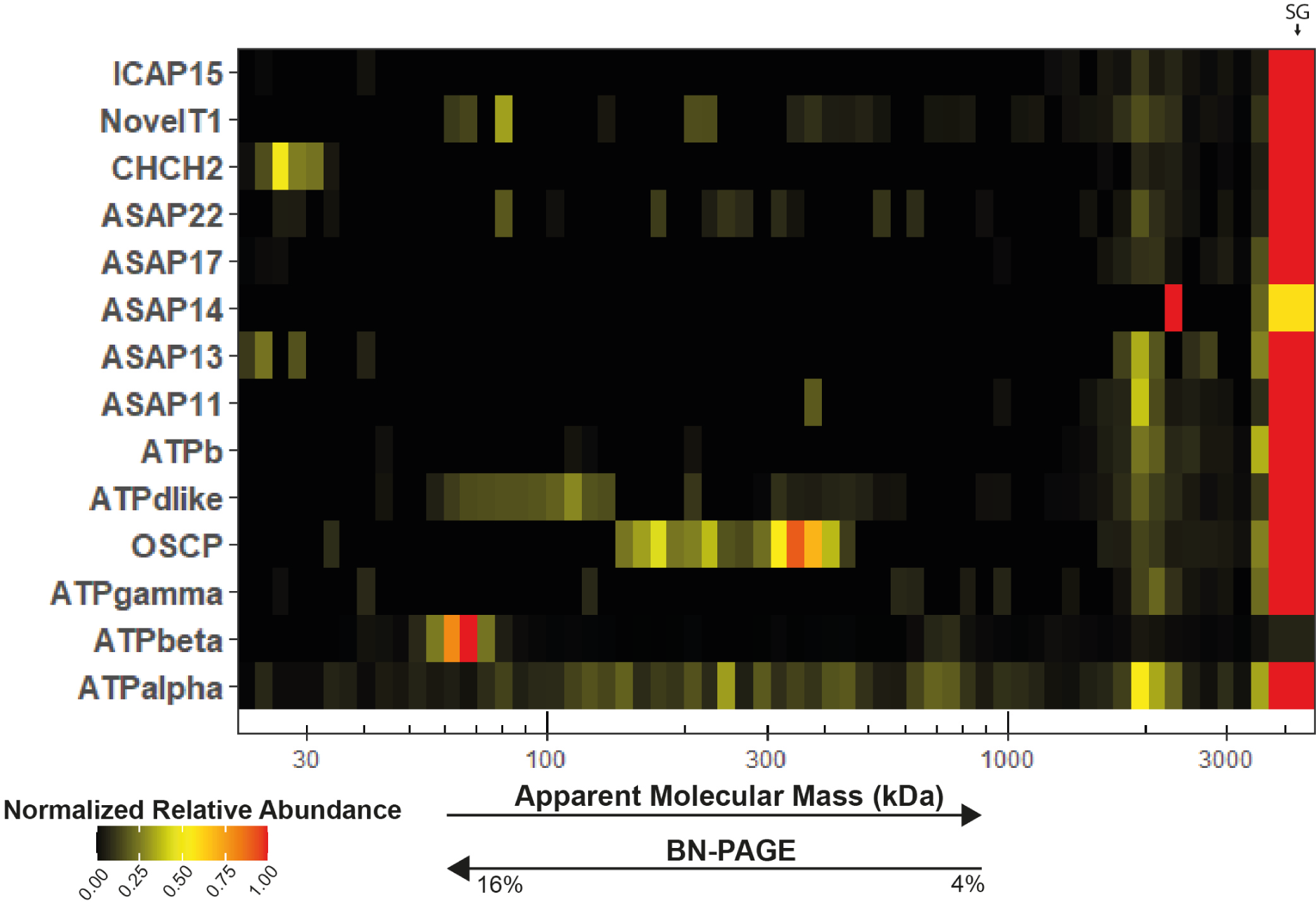
Migration pattern of ATP synthase components in ABS3D. Heatmap showing comigration of putative ATP synthase components in sample ABS3D. Black arrow indicates postion

**Supplementary Figure 7.**
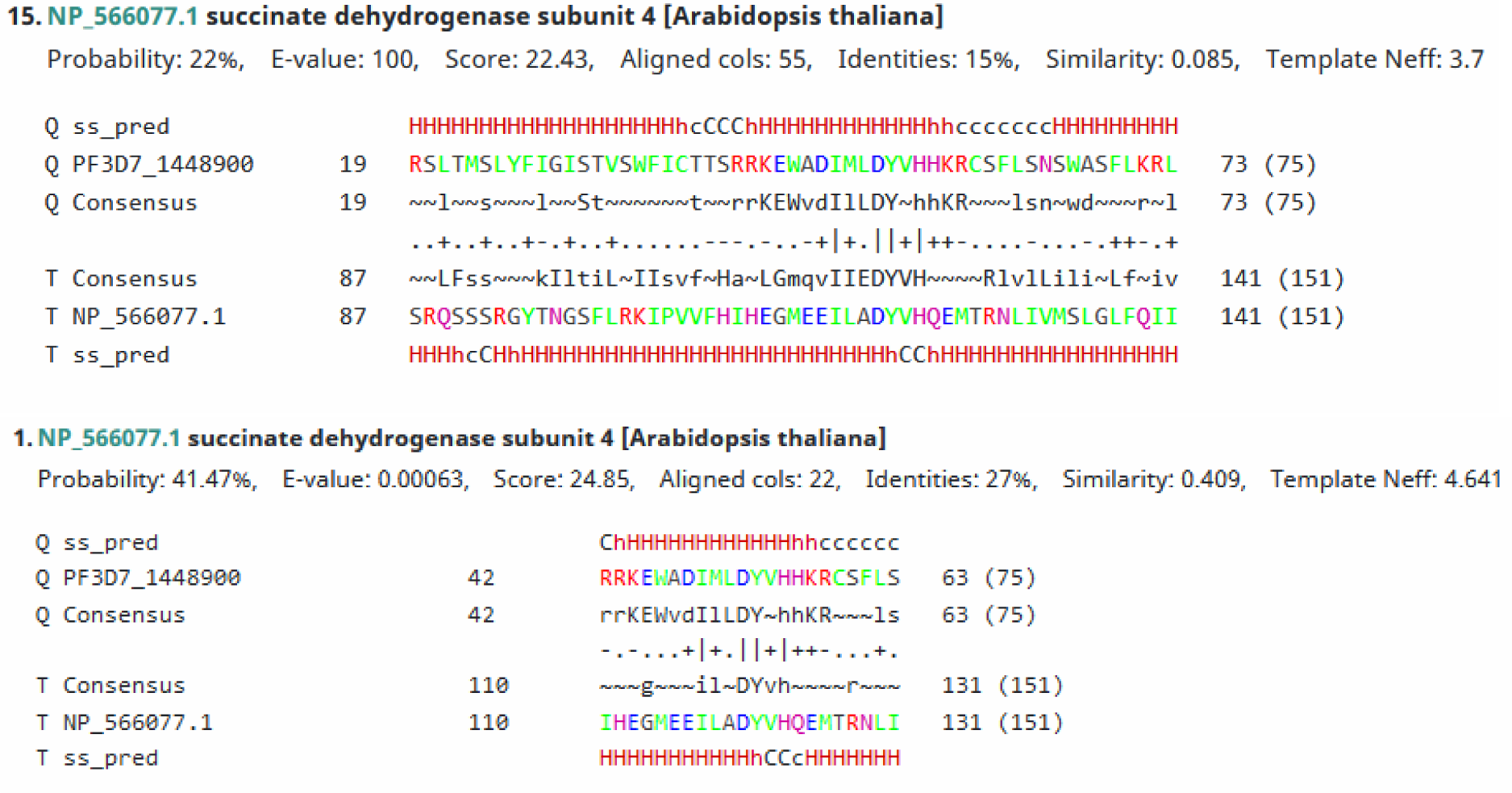
Alignment of PF3D7_1448900 with SDH4 from *A. thaliana*. The alignment was obtained by searching with the PF3D7_1448900 sequence against the profiles of *A. thaliana* using HHpred with default settings (Upper panel). If a pairwise search with SDH4 from A. thaliana is performed, e-value improves and alignment highlights conserved DY motif. The DY motif has been associated with the binding of quinone in other species.

**Supplementary Figure 8.**
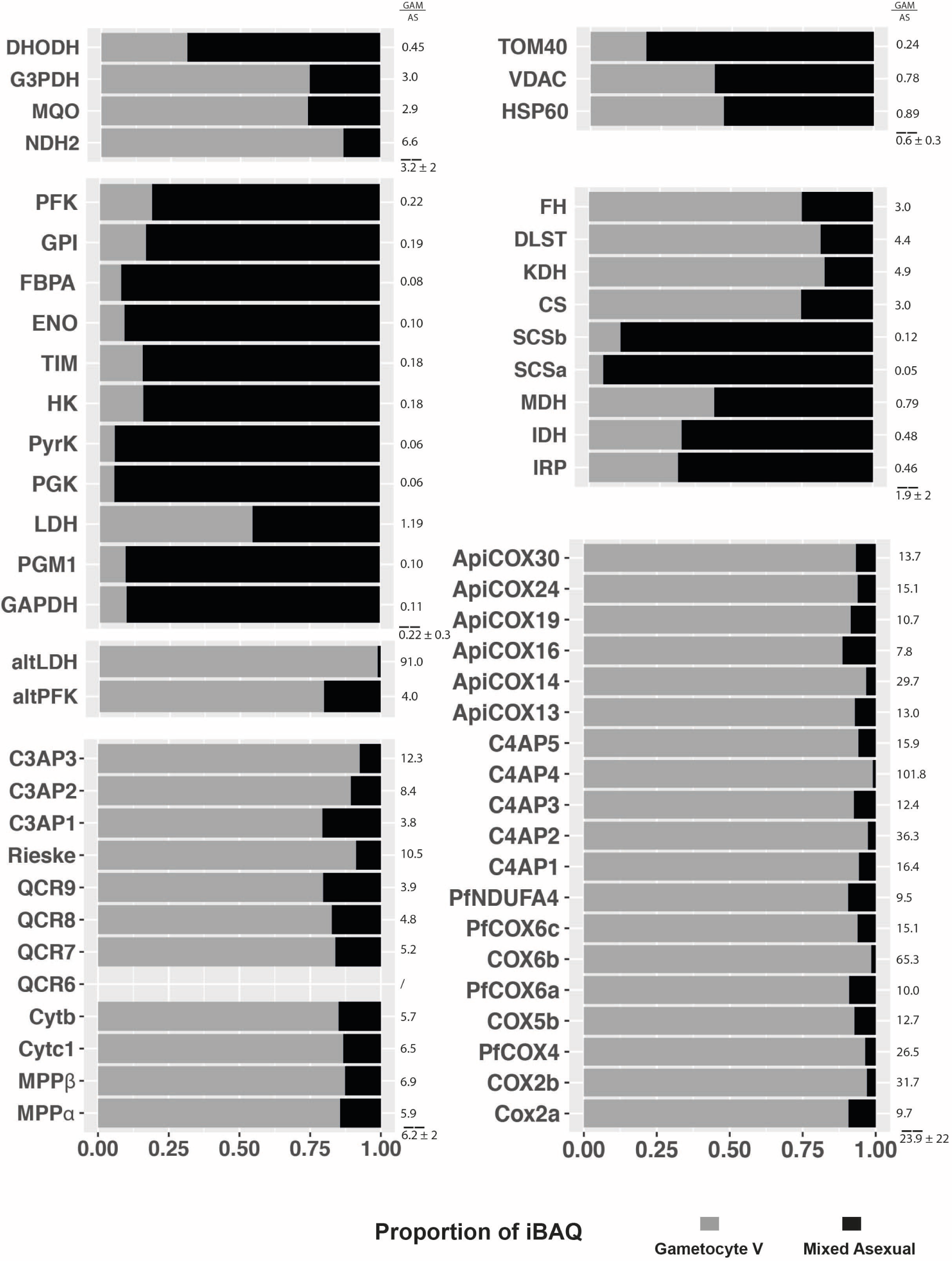
Relative abundance comparisons of respiratory chain complexes III and IV and detected enzymes related to energy metabolism based. Average total iBAQ values of four digitonin ABS parasite and gametocyte complexome samples, respectively, were calculated for individual proteins. VDAC, TOM40, and HSP60 were quantified to approximate degree of mitochondrial enrichment between stages.

## References

1. Vaidya AB, Mather MW. Mitochondrial evolution and functions in malaria parasites. Annu Rev Microbiol. 2009;63:249–67.

2. Vaidya AB, Akella R, Suplick K. Sequences similar to genes for two mitochondrial proteins and portions of ribosomal RNA in tandemly arrayed 6-kilobase-pair DNA of a malarial parasite. Mol Biochem Parasitol. 1989;35(2):97–107.

3. Srivastava IK, Morrisey JM, Darrouzet E, Daldal F, Vaidya AB. Resistance mutations reveal the atovaquone-binding domain of cytochrome b in malaria parasites. Mol Microbiol. 1999;33(4):704–11.

4. Painter HJ, Morrisey JM, Mather MW, Vaidya AB. Specific role of mitochondrial electron transport in blood-stage Plasmodium falciparum. Nature. 2007;446(7131):88–91.

5. Ke H, Lewis IA, Morrisey JM, McLean KJ, Ganesan SM, Painter HJ, et al. Genetic investigation of tricarboxylic acid metabolism during the Plasmodium falciparum life cycle. Cell Rep. 2015;11(1):164–74.

6. Goodman CD, Siregar JE, Mollard V, Vega-Rodriguez J, Syafruddin D, Matsuoka H, et al. Parasites resistant to the antimalarial atovaquone fail to transmit by mosquitoes. Science. 2016;352(6283):349–53.

7. Sturm A, Mollard V, Cozijnsen A, Goodman CD, McFadden GI. Mitochondrial ATP synthase is dispensable in blood-stage Plasmodium berghei rodent malaria but essential in the mosquito phase. Proc Natl Acad Sci U S A. 2015;112(33):10216–23.

8. Matz JM, Goosmann C, Matuschewski K, Kooij TWA. An Unusual Prohibitin Regulates Malaria Parasite Mitochondrial Membrane Potential. Cell Rep. 2018;23(3):756–67.

9. Kanehisa M. Toward understanding the origin and evolution of cellular organisms. Protein Sci. 2019;28(11):1947–51.

10. Seidi A, Muellner-Wong LS, Rajendran E, Tjhin ET, Dagley LF, Aw VY, et al. Elucidating the mitochondrial proteome of Toxoplasma gondii reveals the presence of a divergent cytochrome c oxidase. Elife. 2018;7.

11. Salunke R, Mourier T, Banerjee M, Pain A, Shanmugam D. Correction: Highly diverged novel subunit composition of apicomplexan F-type ATP synthase identified from Toxoplasma gondii. PLoS Biol. 2019;17(3):e3000176.

12. Huet D, Rajendran E, van Dooren GG, Lourido S. Identification of cryptic subunits from an apicomplexan ATP synthase. eLife. 2018;7:e38097.

13. Yu FD, Yang SY, Li YY, Hu W. Co-expression network with protein-protein interaction and transcription regulation in malaria parasite Plasmodium falciparum. Gene. 2013;518(1):7–16.

14. Ramaprasad A, Pain A, Ravasi T. Defining the protein interaction network of human malaria parasite Plasmodium falciparum. Genomics. 2012;99(2):69–75.

15. Lindner SE, Swearingen KE, Shears MJ, Walker MP, Vrana EN, Hart KJ, et al. Transcriptomics and proteomics reveal two waves of translational repression during the maturation of malaria parasite sporozoites. Nat Commun. 2019;10(1):4964.

16. LaCount DJ, Vignali M, Chettier R, Phansalkar A, Bell R, Hesselberth JR, et al. A protein interaction network of the malaria parasite Plasmodium falciparum. Nature. 2005;438(7064):103–7.

17. Rudashevskaya EL, Sickmann A, Markoutsa S. Global profiling of protein complexes: current approaches and their perspective in biomedical research. Expert Review of Proteomics. 2016;13(10):951–64.

18. Wittig I, Braun H-P, Schägger H. Blue native PAGE. Nature protocols. 2006;1(1):418.

19. Heide H, Bleier L, Steger M, Ackermann J, Dröse S, Schwamb B, et al. Complexome profiling identifies TMEM126B as a component of the mitochondrial complex I assembly complex. Cell Metab. 2012;16(4):538–49.

20. Senkler J, Senkler M, Eubel H, Hildebrandt T, Lengwenus C, Schertl P, et al. The mitochondrial complexome of Arabidopsis thaliana. The Plant Journal. 2017;89(6):1079–92.

21. Kahlhöfer F, Kmita K, Wittig I, Zwicker K, Zickermann V. Accessory subunit NUYM (NDUFS4) is required for stability of the electron input module and activity of mitochondrial complex I. Biochimica et Biophysica Acta (BBA) - Bioenergetics. 2017;1858(2):175–81.

22. Hillier C, Pardo M, Yu L, Bushell E, Sanderson T, Metcalf T, et al. Landscape of the Plasmodium Interactome Reveals Both Conserved and Species-Specific Functionality. Cell Reports. 2019;28(6):1635-47.e5.

23. Krungkrai J, Prapunwattana P, Krungkrai SR. Ultrastructure and function of mitochondria in gametocytic stage of Plasmodium falciparum. Parasite. 2000;7(1):19–26.

24. Prince FP. Lamellar and tubular associations of the mitochondrial cristae: unique forms of the cristae present in steroid-producing cells. Mitochondrion. 2002;1(4):381–9.

25. Köhler S. Multi-membrane-bound structures of Apicomplexa: II. the ovoid mitochondrial cytoplasmic (OMC) complex of Toxoplasma gondii tachyzoites. Parasitol Res. 2006;98(4):355–69.

26. Okamoto N, Spurck TP, Goodman CD, McFadden GI. Apicoplast and Mitochondrion in Gametocytogenesis of <em>Plasmodium falciparum</em>. Eukaryotic Cell. 2009;8(1):128–32.

27. Francis G, Kerem Z, Makkar HPS, Becker K. The biological action of saponins in animal systems: a review. British Journal of Nutrition. 2002;88(6):587–605.

28. Filarsky M, Fraschka SA, Niederwieser I, Brancucci NM, Carrington E, Carrió E, et al. GDV1 induces sexual commitment of malaria parasites by antagonizing HP1-dependent gene silencing. Science. 2018;359(6381):1259–63.

29. Wideman JG. The ubiquitous and ancient ER membrane protein complex (EMC): tether or not? F1000Research. 2015;4.

30. Bahl A, Brunk B, Crabtree J, Fraunholz MJ, Gajria B, Grant GR, et al. PlasmoDB: the Plasmodium genome resource. A database integrating experimental and computational data. Nucleic acids research. 2003;31(1):212–5.

31. Ninagawa S, Okada T, Sumitomo Y, Horimoto S, Sugimoto T, Ishikawa T, et al. Forcible destruction of severely misfolded mammalian glycoproteins by the non-glycoprotein ERAD pathway. Journal of Cell Biology. 2015;211(4):775–84.

32. Bai L, You Q, Feng X, Kovach A, Li H. Structure of the ER membrane complex, a transmembrane-domain insertase. Nature. 2020;584(7821):475–8.

33. Kudze T, Mendez-Dorantes C, Jalloh CS, McClellan AJ. Evidence for interaction between Hsp90 and the ER membrane complex. Cell Stress and Chaperones. 2018;23(5):1101–15.

34. Sessler N, Krug K, Nordheim A, Mordmuller B, Macek B. Analysis of the Plasmodium falciparum proteasome using Blue Native PAGE and label-free quantitative mass spectrometry. Amino Acids. 2012;43(3):1119–29.

35. Ito D, Schureck MA, Desai SA. An essential dual-function complex mediates erythrocyte invasion and channel-mediated nutrient uptake in malaria parasites. eLife. 2017;6:e23485.

36. Counihan NA, Chisholm SA, Bullen HE, Srivastava A, Sanders PR, Jonsdottir TK, et al. Plasmodium falciparum parasites deploy RhopH2 into the host erythrocyte to obtain nutrients, grow and replicate. eLife. 2017;6:e23217.

37. Saeed S, Tremp AZ, Dessens JT. The Plasmodium LAP complex affects crystalloid biogenesis and oocyst cell division. International Journal for Parasitology. 2018;48(14):1073–8.

38. Simon N, Scholz SM, Moreira CK, Templeton TJ, Kuehn A, Dude MA, et al. Sexual stage adhesion proteins form multi-protein complexes in the malaria parasite Plasmodium falciparum. J Biol Chem. 2009;284(21):14537–46.

39. Saeed S, Carter V, Tremp AZ, Dessens JT. Translational repression controls temporal expression of the Plasmodium berghei LCCL protein complex. Molecular and Biochemical Parasitology. 2013;189(1):38–42.

40. Lasonder E, Rijpma SR, van Schaijk Ben CL, Hoeijmakers Wieteke AM, Kensche PR, Gresnigt MS, et al. Integrated transcriptomic and proteomic analyses of P. falciparum gametocytes: molecular insight into sex-specific processes and translational repression. Nucleic Acids Research. 2016;44(13):6087–101.

41. Lopez-Barragan MJ, Lemieux J, Quinones M, Williamson KC, Molina-Cruz A, Cui K, et al. Directional gene expression and antisense transcripts in sexual and asexual stages of Plasmodium falciparum. BMC Genomics. 2011;12:587.

42. Scholz SM, Simon N, Lavazec C, Dude MA, Templeton TJ, Pradel G. PfCCp proteins of Plasmodium falciparum: gametocyte-specific expression and role in complement-mediated inhibition of exflagellation. Int J Parasitol. 2008;38(3-4):327–40.

43. de Koning-Ward TF, Gilson PR, Boddey JA, Rug M, Smith BJ, Papenfuss AT, et al. A newly discovered protein export machine in malaria parasites. Nature. 2009;459(7249):945–9.

44. Beck JR, Muralidharan V, Oksman A, Goldberg DE. PTEX component HSP101 mediates export of diverse malaria effectors into host erythrocytes. Nature. 2014;511(7511):592–5.

45. Ho C-M, Beck JR, Lai M, Cui Y, Goldberg DE, Egea PF, et al. Malaria parasite translocon structure and mechanism of effector export. Nature. 2018;561(7721):70–5.

46. Matz JM, Ingmundson A, Costa Nunes J, Stenzel W, Matuschewski K, Kooij TW. In Vivo Function of PTEX88 in Malaria Parasite Sequestration and Virulence. Eukaryot Cell. 2015;14(6):528–34.

47. Garten M, Nasamu AS, Niles JC, Zimmerberg J, Goldberg DE, Beck JR. EXP2 is a nutrient-permeable channel in the vacuolar membrane of Plasmodium and is essential for protein export via PTEX. Nat Microbiol. 2018;3(10):1090–8.

48. Glaser E, Dessi P. Integration of the Mitochondrial-Processing Peptidase into the Cytochrome bc1 Complex in Plants. Journal of Bioenergetics and Biomembranes. 1999;31(3):259–74.

49. Zara V, Conte L, Trumpower BL. Biogenesis of the yeast cytochrome bc1 complex. Biochimica et Biophysica Acta (BBA) - Molecular Cell Research. 2009;1793(1):89–96.

50. Waller RF, Keeling PJ. Alveolate and chlorophycean mitochondrial cox2 genes split twice independently. Gene. 2006;383:33–7.

51. Zimmermann L, Stephens A, Nam SZ, Rau D, Kubler J, Lozajic M, et al. A Completely Reimplemented MPI Bioinformatics Toolkit with a New HHpred Server at its Core. J Mol Biol. 2018;430(15):2237–43.

52. Finn RD, Clements J, Arndt W, Miller BL, Wheeler TJ, Schreiber F, et al. HMMER web server: 2015 update. Nucleic Acids Res. 2015;43(W1):W30–8.

53. Huynen MA, Duarte I, Szklarczyk R. Loss, replacement and gain of proteins at the origin of the mitochondria. Biochim Biophys Acta. 2013;1827(2):224–31.

54. Hartley AM, Lukoyanova N, Zhang Y, Cabrera-Orefice A, Arnold S, Meunier B, et al. Structure of yeast cytochrome c oxidase in a supercomplex with cytochrome bc(1). Nat Struct Mol Biol. 2019;26(1):78–83.

55. Zhu G, Marchewka MJ, Keithly JS. Cryptosporidium parvum appears to lack a plastid genome. Microbiology. 2000;146(2):315–21.

56. Lipper CH, Karmi O, Sohn YS, Darash-Yahana M, Lammert H, Song L, et al. Structure of the human monomeric NEET protein MiNT and its role in regulating iron and reactive oxygen species in cancer cells. Proceedings of the National Academy of Sciences. 2018;115(2):272–7.

57. Salunke R, Mourier T, Banerjee M, Pain A, Shanmugam D. Correction: Highly diverged novel subunit composition of apicomplexan F-type ATP synthase identified from Toxoplasma gondii. PLoS biology. 2019;17(3):e3000176–e.

58. Orczyk M, Wojciechowski K, Brezesinski G. The influence of steroidal and triterpenoid saponins on monolayer models of the outer leaflets of human erythrocytes, E. coli and S. cerevisiae cell membranes. Journal of Colloid and Interface Science. 2020;563:207–17.

59. Balabaskaran Nina P, Morrisey JM, Ganesan SM, Ke H, Pershing AM, Mather MW, et al. ATP synthase complex of Plasmodium falciparum: dimeric assembly in mitochondrial membranes and resistance to genetic disruption. J Biol Chem. 2011;286(48):41312–22.

60. Tanaka TQ, Hirai M, Watanabe Y-i, Kita K. Toward understanding the role of mitochondrial complex II in the intraerythrocytic stages of Plasmodium falciparum: Gene targeting of the Fp subunit. Parasitology International. 2012;61(4):726–8.

61. Mogi T, Kita K. Identification of mitochondrial Complex II subunits SDH3 and SDH4 and ATP synthase subunits a and b in Plasmodium spp. Mitochondrion. 2009;9(6):443–53.

62. Morales J, Mogi T, Mineki S, Takashima E, Mineki R, Hirawake H, et al. Novel mitochondrial complex II isolated from Trypanosoma cruzi is composed of 12 peptides including a heterodimeric Ip subunit. The Journal of biological chemistry. 2009;284(11):7255–63.

63. Silkin Y, Oyedotun KS, Lemire BD. The role of Sdh4p Tyr-89 in ubiquinone reduction by the Saccharomyces cerevisiae succinate dehydrogenase. Biochim Biophys Acta. 2007;1767(2):143–50.

64. Klug D, Mair GR, Frischknecht F, Douglas RG. A small mitochondrial protein present in myzozoans is essential for malaria transmission. Open Biol. 2016;6(4):160034.

65. Kurokawa T, Sakamoto J. Purification and characterization of succinate: menaquinone oxidoreductase from Corynebacterium glutamicum. Archives of microbiology. 2005;183(5):317–24.

66. Yankovskaya V, Horsefield R, Tornroth S, Luna-Chavez C, Miyoshi H, Leger C, et al. Architecture of succinate dehydrogenase and reactive oxygen species generation. Science. 2003;299(5607):700–4.

67. Salunke R, Mourier T, Banerjee M, Pain A, Shanmugam D. Highly diverged novel subunit composition of apicomplexan F-type ATP synthase identified from Toxoplasma gondii. PLoS Biol. 2018;16(7):e2006128.

68. Zhang M, Wang C, Otto TD, Oberstaller J, Liao X, Adapa SR, et al. Uncovering the essential genes of the human malaria parasite Plasmodium falciparum by saturation mutagenesis. Science. 2018;360(6388).

69. Sidik SM, Huet D, Ganesan SM, Huynh M-H, Wang T, Nasamu AS, et al. A Genome-wide CRISPR Screen in Toxoplasma Identifies Essential Apicomplexan Genes. Cell. 2016;166(6):1423-35.e12.

70. Barylyuk K, Koreny L, Ke H, Butterworth S, Crook OM, Lassadi I, et al. A subcellular atlas of Toxoplasma reveals the functional context of the proteome. bioRxiv. 2020.

71. MacRae JI, Dixon MW, Dearnley MK, Chua HH, Chambers JM, Kenny S, et al. Mitochondrial metabolism of sexual and asexual blood stages of the malaria parasite Plasmodium falciparum. BMC biology. 2013;11(1):1–10.

72. Srivastava A, Philip N, Hughes KR, Georgiou K, MacRae JI, Barrett MP, et al. Stage-Specific Changes in Plasmodium Metabolism Required for Differentiation and Adaptation to Different Host and Vector Environments. PLOS Pathogens. 2016;12(12):e1006094.

73. Meerstein-Kessel L, van der Lee R, Stone W, Lanke K, Baker DA, Alano P, et al. Probabilistic data integration identifies reliable gametocyte-specific proteins and transcripts in malaria parasites. Sci Rep. 2018;8(1):410.

74. Cogliati S, Enriquez JA, Scorrano L. Mitochondrial Cristae: Where Beauty Meets Functionality. Trends in Biochemical Sciences. 2016;41(3):261–73.

75. Mony BM, Mehta M, Jarori GK, Sharma S. Plant-like phosphofructokinase from Plasmodium falciparum belongs to a novel class of ATP-dependent enzymes. International Journal for Parasitology. 2009;39(13):1441–53.

76. van Esveld SL, Huynen MA. Does mitochondrial DNA evolution in metazoa drive the origin of new mitochondrial proteins? IUBMB Life. 2018;70(12):1240–50.

77. Lhouvum K, Balaji S, Ahsan MJ, Trivedi V. Plasmodium falciparum PFI1625c offers an opportunity to design potent anti-malarials: Biochemical characterization and testing potentials in drug discovery. Acta tropica. 2019;191:116–27.

78. Lhouvum K, Bhuyar KS, Trivedi V. Molecular modeling and correlation of PFI1625c-peptide models of bioactive peptides with antimalarial properties. Medicinal Chemistry Research. 2015;24(4):1527–33.

79. Escalante AA, Ayala FJ. Evolutionary origin of Plasmodium and other Apicomplexa based on rRNA genes. Proceedings of the National Academy of Sciences. 1995;92(13):5793–7.

80. van der Sluis EO, Bauerschmitt H, Becker T, Mielke T, Frauenfeld J, Berninghausen O, et al. Parallel Structural Evolution of Mitochondrial Ribosomes and OXPHOS Complexes. Genome Biol Evol. 2015;7(5):1235–51.

81. Shen Y-Y, Shi P, Sun Y-B, Zhang Y-P. Relaxation of selective constraints on avian mitochondrial DNA following the degeneration of flight ability. Genome Res. 2009;19(10):1760–5.

82. Blum TB, Hahn A, Meier T, Davies KM, Kühlbrandt W. Dimers of mitochondrial ATP synthase induce membrane curvature and self-assemble into rows. Proceedings of the National Academy of Sciences. 2019;116(10):4250–5.

83. Mühleip AW, Joos F, Wigge C, Frangakis AS, Kühlbrandt W, Davies KM. Helical arrays of U-shaped ATP synthase dimers form tubular cristae in ciliate mitochondria. Proceedings of the National Academy of Sciences. 2016;113(30):8442–7.

84. Trager W, Jensen JB. Human malaria parasites in continuous culture. Science. 1976;193(4254):673–5.

85. Lambros C, Vanderberg JP. Synchronization of Plasmodium falciparum erythrocytic stages in culture. The Journal of parasitology. 1979:418–20.

86. Fivelman QL, McRobert L, Sharp S, Taylor CJ, Saeed M, Swales CA, et al. Improved synchronous production of Plasmodium falciparum gametocytes in vitro. Molecular and Biochemical Parasitology. 2007;154(1):119–23.

87. Ribaut C, Berry A, Chevalley S, Reybier K, Morlais I, Parzy D, et al. Concentration and purification by magnetic separation of the erythrocytic stages of all human Plasmodium species. Malaria Journal. 2008;7(1):45.

88. Heide H, Bleier L, Steger M, Ackermann J, Dröse S, Schwamb B, et al. Complexome profiling identifies TMEM126B as a component of the mitochondrial complex I assembly complex. Cell metabolism. 2012;16(4):538–49.

89. Olsen JV, de Godoy LM, Li G, Macek B, Mortensen P, Pesch R, et al. Parts per million mass accuracy on an Orbitrap mass spectrometer via lock mass injection into a C-trap. Molecular & Cellular Proteomics. 2005;4(12):2010–21.

90. Tyanova S, Temu T, Cox J. The MaxQuant computational platform for mass spectrometry-based shotgun proteomics. Nature Protocols. 2016;11(12):2301–19.

91. de Hoon MJL, Imoto S, Nolan J, Miyano S. Open source clustering software. Bioinformatics. 2004;20(9):1453–4.

92. Team RC. R: A language and environment for statistical computing. Vienna, Austria; 2013.

93. Wickham H. ggplot2. WIREs Computational Statistics. 2011;3(2):180–5.

94. Potter SC, Luciani A, Eddy SR, Park Y, Lopez R, Finn RD. HMMER web server: 2018 update. Nucleic Acids Research. 2018;46(W1):W200–W4.

95. Hartley AM, Meunier B, Pinotsis N, Marechal A. Rcf2 revealed in cryo-EM structures of hypoxic isoforms of mature mitochondrial III-IV supercomplexes. Proc Natl Acad Sci U S A. 2020;117(17):9329–37.

96. Krieger E, Vriend G. YASARA View - molecular graphics for all devices - from smartphones to workstations. Bioinformatics. 2014;30(20):2981–2.

